# Horizontally transferred genes as RNA interference targets for aphid and whitefly control

**DOI:** 10.1101/2022.10.07.511359

**Authors:** Honglin Feng, Wenbo Chen, Sonia Hussain, Sara Shakir, Vered Tzin, Femi Adegbayi, Todd Ugine, Zhangjun Fei, Georg Jander

## Abstract

RNA interference (RNAi)-based technologies are starting to be commercialized as a new approach for agricultural pest control. Horizontally transferred genes (HTGs), which have been transferred into insect genomes from viruses, bacteria, fungi, or plants, are attractive targets for RNAi-mediated pest control. HTGs are often unique to a specific insect family or even genus, making it unlikely that RNAi constructs targeting such genes will have negative effects on ladybugs, lacewings, and other beneficial predatory insect species. In this study, we sequenced the genome of a red, tobacco-adapted isolate of *Myzus persicae* (green peach aphid) and bioinformaticaly identified 30 HTGs. We then used plant-mediated virus-induced gene silencing (VIGS) to show that several HTGs of bacterial and plant origin are important for aphid growth and/or survival. Silencing the expression of fungal HTGs did not affect aphid survivorship, but decreased aphid reproduction. Importantly, although there was uptake of plant-expressed RNA by *Coccinella septempunctata* (seven-spotted ladybugs) via the aphids that they consumed, we did not observe negative effects on ladybugs from aphid-targeted VIGS constructs. In other experiments, we targeted five *Bemisia tabaci* (whitefly) HTGs using VIGS and demonstrated that knockdown of some of these genes affected whitefly survival. As functional HTGs have been identified in the genomes of numerous pest species, we propose that these HTGs should be explored further as efficient and safe targets for control of insect pests using plant-mediated RNA interference.

## Introduction

Aphids, whiteflies, and other hemipteran insect pests cause considerable damage to agricultural crops. Although current control strategies, in particular chemical pesticides, provide some level of protection, insects continuously develop new tolerance or resistance mechanisms (Bass et al., 2014; Koch et al., 2018). Genetically engineered *Bt* crops have been shown to be effective in controlling specific insect pests, with limited environmental persistence (Sears et al., 2001; Mendelsohn et al., 2003; Sujii et al., 2013; Tian et al., 2015). However, there has been only limited success in the development of *Bt* toxins that target hemipteran pests (Lu et al., 2010; Chougule and Bonning, 2012; Sujii et al., 2013; Pessoa et al., 2016). Thus, there is a need to develop novel and environmentally friendly transgenic plant approaches for the control of phloem-feeding insects such as aphids and whiteflies.

RNA interference (RNAi)-based gene expression silencing has emerged as a novel and powerful strategy for agricultural pest control. As phloem feeders, hemipteran pests are less likely to take up surface sprays of RNAi constructs, which can be deployed against chewing herbivores such as Colorado potato beetles (*Leptinotarsa decemlineata*) (Rodrigues et al., 2021). However, plant-mediated RNAi has been effective for targeting hemipteran gene expression (Chung et al., 2021). Several classes of aphid genes have been successfully inhibited by plant-mediated RNAi, including *C002* (Mutti et al., 2006; Pitino et al., 2011; Coleman et al., 2015), receptor of activated kinase C (*Rack-1*) (Pitino et al., 2011; Coleman et al., 2015), *MpPInt01* and *MpPInt02* (Coleman et al., 2015), *Mp55* (Elzinga et al., 2014); aquaporin, sucrase-transglucosidase, and sugar transporters (Tzin et al., 2015), V-ATPase E and tubulin folding cofactor D (Guo et al., 2014), serine protease (Bhatia et al., 2012), hunchback (Mao and Zeng, 2014), carboxylesterase CbE E4, juvenile hormone binding protein, calreticulin, and cathepsin, carboxylesterase (Xu et al., 2014), salivary sheath protein (Abdellatef et al., 2015), and a zinc finger protein gene (Xie et al., 2022). Similarly, genes that have been effectively targeted using plant-mediated RNAi for controlling *Bemisia tabaci* (whiteflies) include ecdysone receptor (EcR) and acetylcholinesterase (AchE) (Malik et al., 2016), cytochrome P450 genes cyp315a1 and cyp18a1(Luan et al., 2013), aquaporin and α-glucosidase (Raza et al., 2016), v-ATPase (Thakur et al., 2014; Ibrahim et al., 2017), cyclophilin B (CypB) and heat shock protein 70 (hsp70) (Kanakala et al., 2019), and phenolic glucoside malonlyltransferase (*BtPMaT1*) (Xia et al., 2021).

As part of the regulatory approval process for commercialization, RNAi-based transgenic plants should be assessed for biological risks, which include effects on non-target insect species with similar gene sequences (Casacuberta et al., 2015). Essential genes that are potential targets for RNAi-mediated pest control, including many of those described above for the control of aphids and whiteflies, are often highly conserved across the insect phylogeny. As it is not currently feasible to examine the genomes of all potential non-target insects, bioinformatic analysis alone is not sufficient for the design of RNAi constructs that are unlikely to have off-target effects on beneficial insect species.

Horizontally transferred genes (HTGs), which are often taxon-, genus-, or even species-specific (Wybouw et al., 2016), are an attractive option for limiting potential off-target effects during RNAi-mediated pest control. HTGs, including transfers from prokaryote to prokaryote, prokaryote to eukaryote, and eukaryote to eukaryote, have been described in all branches of life (Soucy et al., 2015; Husnik and McCutcheon, 2018). Importantly, the integration, expression, and maintenance HTGs in the recipient genome suggests that their presence provides a selective advantage (Soucy et al., 2015). There are many hurdles for HTGs to become functional in the recipient genome, particularly in the case of genes that are transformed from prokaryotes to eukaryotes. For instance, the presence or absence of introns, variable GC content, codon usage preferences, and differences in transcriptional promoters can limit successful gene expression in a recipient species (Husnik and McCutcheon, 2018). Nevertheless, functional HTGs have been described in many species, including aphids and whiteflies (Novakova and Moran, 2012; Sloan et al., 2014; Luan et al., 2015; Chen et al., 2016; Chung et al., 2018; Parker and Brisson, 2019).

Prior to this study, a few horizontally transferred genes with different origins has been described in the aphid genomes. Carotenoid biosynthetic genes in aphids were shown to be horizontally transferred from a fungus, including the highly duplicated carotene desaturase (*Tor*) and the carotenoid cyclase-carotenoid synthase (*CarRP*) (Novakova and Moran, 2012). One carotenoid desaturase gene copy was present only in a red *Myzus persicae* (green peach aphid) isolate, and a knockout mutation resulted in the loss of red color in the aphids (Moran and Jarvik, 2010). Similar red color reduction was also observed in *Acyrthosiphon pisum* (pea aphid) nymphs when silencing a carotenoid desaturase in the parental aphids (Ding et al., 2020). These results indicated that fungal-origin carotenoid biosynthetic genes remain functional in the recipient aphid genomes and confer to the aphid red-green color polymorphism, in turn influence their susceptibility to natural enemies (Moran and Jarvik, 2010). More recently, two *A. pisum* (pea aphid) HTGs of bacterial origin, *amiD* and *ldcA1*, were targeted using RNAi constructs that were supplied in artificial diet (Chung et al., 2018). Enriched *amidD* and *ldcA1* expression in *A. pisum* bacteriocytes, which harbor *Buchnera aphidicola* endosymbionts, coupled with the fact that *amiD* and *ldcA* were lost when the peptidoglycan biosynthetic genes were not present in the symbiont *Buchnera* genome (Smith et al., 2022), suggested that the amiD and ldcA1 function in degrading bacterial peptidoglycan, thereby protecting *Buchnera* from host attack (Chung et al., 2018). Consistent with this hypothesis, knockdown of *amiD* and *ldcA1* by RNAi caused a significant reduction in *Buchnera* abundance and inhibited aphid growth. In addition to HTGs from fungi and bacterial, HTGs originated from virus have also been found in aphids. For example, the cytolethal distending toxin subunit B (*cdtB*) found in the genome of *M. persicae* strain G006 was suggested to be involved in aphid resistance to a predatory wasp (Verster et al., 2019). In the pea aphid genome, two horizontally transferred genes from a densovirus have been described and demonstrated to modulate the aphid wing plasticity (Parker and Brisson, 2019).

Our previous analysis of the *B. tabaci* MEAM1 genome identified 142 HTGs from bacteria, fungi, and plants (Chen et al., 2016). Several of these genes were proposed to contribute to broad host range and insecticide resistance of whiteflies. Interestingly, a recent study found that a detoxifying gene in whiteflies, phenolic glucoside malonyltransferase (*BtPMaT1*), likely originated from plants and was able to neutralize host plant defensive metabolites (Xia et al., 2021). Plant-mediated silencing of *BtPMaT1* expression impairs whitefly detoxification functions and confers full whitefly resistance in tomato plants, confirming the utility of this HTG for whitefly control (Xia et al., 2021).

To further explore HTGs as potential RNAi targets for aphid control, we assembled genome of a red, tobacco-adapted strain of *M. persicae* (Ramsey et al., 2007; Ramsey et al., 2014), conducted a genome-wide annotation of HTGs, and compared this to the HTG profiles of other aphid species. Then, using plant-mediated virus-induced gene silencing (VIGS), we tested the RNAi effects of these HTGs on aphid survival. Additionally, we conducted RNAi of a subset of the HTGs that have been annotated in the *B. tabaci* MEAM1 (whitefly) (Chen et al., 2016).

We found that knock-down of these HTGs using plant-mediated RNAi can significantly affect insect performance. Importantly, the knock-down of the aphid HTGs did not affect the survival of *Coccinella septempunctata* (seven-spotted ladybugs) that fed on these aphids. Our results suggest that HTGs will be effective and safe targets for plant genetic engineering to control aphid populations in the field.

## Materials and methods

### Insect and plant cultures

The tobacco-adapted *M. persicae* strain USDA-Red (Ramsey et al., 2007; Ramsey et al., 2014) was maintained on *Nicotiana tabacum* (tobacco) plants in a growth room at 23°C with a 16:8 h light:dark photoperiod. A *B. tabaci* MEAM1 colony was obtained from Angela Douglas (Cornell University) and maintained on an acylsugar-deficient *asat2-1* mutant *Nicotiana benthamiana* lineage (Feng et al., 2021) at 23°C with a 16:8 h light:dark photoperiod. A *C. septempunctata* colony was maintained on *Acyrthosiphon pisum* (pea aphid) feeding on *Vicia faba* (faba bean). Mealybug ladybird beetle (*Cryptolaemus montrouzieri*) adults were purchased from Amazon (www.amazon.com). *Nicotiana benthamiana* wild type and *asat2-1* mutant plants (Feng et al., 2021) for aphid, whitefly, and ladybug experiments were grown in Cornell Mix (by weight 56% peat moss, 35% vermiculite, 4% lime, 4% Osmocoat slow-release fertilizer [Scotts, Marysville, OH], and 1% Unimix [Scotts]) in a Conviron growth chamber with a photosynthetic photon flux density of 200 mmol m-1 s-1 and a 16:8 h day:night photoperiod, at 23°C with 50% relative humidity.

### Long-read PacBio sequencing

To sequence the USDA-Red *M. persicae* genome, we extracted high molecular weight genomic DNA (gDNA) from 100–200 mg of fresh mixed-instar insect tissues using a protocol described previously (Fulton et al., 1995; Chen et al., 2019). Briefly, whole aphids collected from *N. tabacum* plants were ground in liquid nitrogen, and mixed with 400 µl microprep buffer made up of DNA extraction buffer (0.35 M sorbitol, 0.1 M Tris-base, 5 mM EDTA [ethylenediaminetetraacetic acid], pH 7.5.), nuclei lysis buffer (0.2 M Tris-base pH 7.5, 0.05 M EDTA, 2 M NaC1, 2 % CTAB [cetyltrimethylammonium bromide]), 5% Sarkosyl, and freshly added 0.5% sodium bisulfite, followed by incubation at 65°C for 30 minutes. Next, 400 µl chloroform:isoamyl alcohol (24:1) was added to the cooled-down sample, mixed vigorously, and centrifuged at 23°C for 10 min at 14,000 × g. The upper aqueous phase was transferred to a new 1.5 ml microcentrifuge tube with 3 µl of RNase A to remove RNA and incubated at 37°C for 15 min, followed by repeated washing with 400 µl chloroform:isoamyl alcohol (24:1), as described above. The supernatant was transferred to a new microcentrifuge tube with 200 µl ice-cold isopropanol and mixed gently by inversion. The DNA was precipitated by centrifugation at 14,000 × g at 4°C for 10 min. The DNA pallet was washed with 70% ethanol, air-dried and dissolved in 50 µl of nuclease-free water. The quantity and quality of purified gDNA was analyzed with Qubit 3 fluorometer (Thermo Fisher, Waltham, MA, USA) and a Bioanalyzer DNA12000 kit (Agilent, Santa Clara, CA, USA), respectively.

Single-molecule real-time sequencing (SMRT) 20 kb DNA template was prepared using the PacBio Sequel 2.0 sequencing enzyme and chemistry according to manufacturer’s (PacBio, Menlo Park, CA, USA) instructions. Briefly, to remove small fragments and/or organic contaminants, approximately 20 μg of DNA was re-purified by adding 0.5 × AMPure XP beads (Beckman Coulter, Indianapolis, IN, USA) and vortexed at 2,000 × g for 10 min, followed by washing twice with 70% ethanol, and DNA was dissolved in the elution buffer. DNA was sheared to 25–30 kb using a Covaris G-tube (Wobern, MA, USA) and an Eppendorf 5424 centrifuge (Hamburg, Germany) at 3,000 g. The DNA was then purified with 0.5 × AMPure XP beads and the average DNA fragment size was determined by Agilent Bioanalyzer DNA12000 kit. The purified DNA was processed through DNA damage and end-repair steps. Briefly, to repair the DNA damage, the fragmented DNA was first treated with 0.18 U/μl of P6 polymerase followed by 52 µl of DNA damage repair solution (1 × DNA damage repeat buffer, 1 × nicotinamide adenine dinucleotide, 1 mM adenosine triphosphate [ATP] high, 0.1 mM dNTP, and 1 × DNA damage repeat mix) and incubated at 37°C for 20 min. To repair DNA ends, 1 × end repair mix was added to the solution and incubated at 25°C for 10 min, followed by another 0.45 × Ampure XP purification step.

To prepare SMRTbell (PacBio) library, 0.75 μM of blunt adapters were first added to DNA followed by adding 1 × template preparation buffer, 0.05 mM ATP low, and 0.75 U/μl T4 ligase in a final volume of 40 μl and incubation at 25°C overnight. Ligase enzyme was denatured at 65°C for 10 min. The unligated fragments were removed by incubating the DNA library at 37°C with an exonuclease cocktail solution made up of 1.81 U/μl Exonuclease III and 0.18 U/μl Exonuclease VII for 1 h, followed by removal of <1,000-bp molecular-weight DNA and biological contaminants by two 0.45 × Ampure XP purification steps. Next, size selection of 17– 50 kb was performed on Sage BluePippin (Sage Science, Beverly, MA, USA) using manufacturer’s instructions. An additional DNA damage repair step was performed on size selected DNA, followed by 0.8 × AMP bead purification. The fragment sizes were verified by Agilent bioanalyzer DNA12000 kit, and the mass was quantified using an Invitrogen Qubit 3 fluorometer (Thermo Fisher). Next, the sequencing reaction was prepared by binding Polymerase 2.0 with DNA template at 30°C for 4 hours. The binding complex was diluted and diffusion-loaded on the plate at 6 pM on a Sequel 5.0 system and sequenced on a Sequel machine with 2.0 chemistry recording 10-h movies at the Icahn Institute and Department of Genetics and Genomic Sciences, Icahn School of Medicine at Mount Sinai (New York, NY, USA). Twelve SMRT cells were run on the PacBio Sequel platform, yielding ∼62 gigabases (Gb) raw sequence data (Table 1) for the *M. persicae* genome. The raw data analysis was performed on SMRTLink 5.0.

**Table 1.** Horizontally transferred genes identified in aphid genomes

| ID in *M. persicae* USDA-Red | Annotation | Origin | *M. persicae* G006 | *A. pisum* | *A. glycines* | *A. gossypii* | *D. vitifoliae* | *D. noxia* | *E. lanigerum* | *M. cerasi* | *P. nigronervosa* | *R. maidis* | *R. padi* |
| --- | --- | --- | --- | --- | --- | --- | --- | --- | --- | --- | --- | --- | --- |
| MPE127429 | Putative LD carboxypeptidase | Bacteria | y | y | y | y | n | y | y | y | y | y | y |
| MPE127428 | ATP binding protein | Bacteria | y | y | y | y | y | y | y | y | y | y | y |
| MPE015894 | N-acetylmuramoyl-L-alanine amidase | Bacteria | y | y | n | n | n | y | n | y | n | n | n |
| MPE005086 | Cytolethal distending toxin subunit B | Viruses | y | n | n | n | n | n | n | y | n | n | n |
| MPE025920 | Non-structural protein NS1 superfamily | Viruses | y | y | y | n | y | n | n | y | y | n | y |
| MPE014463 | Non-structural protein NS1 superfamily | Viruess | y | n | n | n | n | n | n | n | n | n | n |
| MPE006690 | Non-structural protein NS1 superfamily | Viruses | y | n | n | n | n | n | n | n | n | n | n |
| MPE006487 | Lycopene cyclase phytoene synthase | Fungi | y | y | y | y | y | y | y | y | y | y | y |
| MPE006492 | Lycopene cyclase phytoene synthase | Fungi | y | y | y | y | y | y | y | y | y | y | y |
| MPE006495 | Lycopene cyclase phytoene synthase | Fungi | y | y | y | y | y | y | y | y | y | y | y |
| MPE006496 | Lycopene cyclase phytoene synthase | Fungi | y | y | y | y | y | y | y | y | y | y | y |
| MPE006497 | Lycopene cyclase phytoene synthase | Fungi | y | y | y | y | y | y | y | y | y | y | y |
| MPE006498 | Lycopene cyclase phytoene synthase | Fungi | y | y | y | y | y | y | y | y | y | y | y |
| MPE006500 | Lycopene cyclase phytoene synthase | Fungi | y | y | y | y | y | y | y | y | y | y | y |
| MPE006501 | Lycopene cyclase phytoene synthase | Fungi | y | y | y | y | y | y | y | y | y | y | y |
| MPE006504 | Lycopene cyclase phytoene synthase | Fungi | y | y | y | y | y | y | y | y | y | y | y |
| MPE006493 | Carotenoid desaturase | Fungi | y | y | y | y | y | y | y | y | y | y | n |
| MPE006499 | Carotenoid desaturase | Fungi | y | y | y | y | y | y | y | y | y | y | n |
| MPE006502 | Carotenoid desaturase | Fungi | y | y | y | y | y | y | y | y | y | y | n |
| MPE006505 | Carotenoid desaturase | Fungi | y | y | y | y | y | y | y | y | y | y | y |
| MPE002740 | Unknown with ankyrin repeats | Plants | y | y | y | y | n | n | y | y | y | y | y |
| MPE008575 | Unknown with ankyrin repeats | Plants | y | n | n | n | n | n | n | y | n | n | n |
| MPE008576 | Unknown with ankyrin repeats | Planst | y | n | n | n | n | n | n | y | n | n | n |
| MPE010373 | Unknown with ankyrin repeats | Plants | y | y | n | n | n | n | n | y | y | y | y |
| MPE011636 | Reverse transcriptase domain-containing protein | Plants | y | n | n | n | n | n | n | y | n | n | y |
| MPE013670 | Unknown with ankyrin repeats | Plants | y | n | n | n | n | n | n | y | n | n | n |
| MPE016359 | Unknown with ankyrin repeats | Plants | y | y | y | y | n | y | n | y | y | y | y |
| MPE018635 | Unknown with ankyrin repeats | Plants | y | n | n | n | n | n | n | n | n | n | n |
| MPE018636 | Unknown with ankyrin repeats | Plants | y | y | n | n | n | n | n | n | n | n | y |
| MPE023973 | Unknown with ankyrin repeats | Plants | y | n | y | n | n | y | n | y | n | y | y |

### Short-read Illumina sequencing

One paired-end library was generated using Illumina TruSeq DNA sample preparation kit (Illumina, San Diego, CA, USA), following the manufacturer’s instructions. Briefly, DNA quality was first checked with Invitrogen Qubit 3 fluorometer, and ∼2 μg of DNA was dissolved in resuspension buffer in a volume of 55 μl, vortexed for 2 min and centrifuged at 280 × g for 1 min. The DNA sample (52.5 μl) was transferred to a Covaris microTUBE and fragmented by centrifuging at 280 × g for 5 s. Sample purification beads (SPBs) were used to purify the DNA, following by washing twice with 80% ethanol and DNA was then dissolved in 50 μl of resuspension buffer. DNA was end-repaired by incubating at 30°C in 40 μl of end repair mixture for 30 min. The DNA library of insert size ∼550 bp was purified using SPBs according to manufacturer’s instructions. Briefly, larger fragments were removed by adding 160 μl of SPBs solution (50% SPBs in PCR [polymerase chain reaction] grade water), vortexed at 1,800 × g for 2 min, incubated at 23°C for 5 min, and centrifuged again at 280 × g for 1 min. Two hundred fifty μl of supernatant were transferred to a cleanup end repair plate. Small fragments were removed by adding 30 μl of SPBs to the supernatant, vortexed at 1,800 × g for 2 min, incubated at 23°C for 5 min, and centrifuged at 280 × g for 1 min. The SPBs were then washed twice with 80% ethanol, and DNA was dissolved in 15 μl of resuspension buffer. To adenylate the DNA at 3’ end, 2.5 μl of A-Tailing control and 12.5 μl of A-Tailing mixture (New England Biolabs, Ipswich, MA, USA) was added to the DNA solution in a final volume of 30 μl and vortexed at 1,800 × g for 2 min. The solution was first incubated at 37°C for 30 min, then at 70°C for 5 min, and a final incubation on ice for 5 min.

To prepare a TruSeq library, DNA was mixed with 2.5 μl of adapters, 2.5 μl of ligation mixture and 2.5 μl of ligation control and incubated at 30°C for 10 min. The ligation step was stopped by adding 5 μl of ligation stop buffer and ligated fragments were purified using SPBs and washed twice with 80% ethanol. The library was quantified with the KAPA Library Quantification Kit (Roche, Basel, Switzerland), and the fragment size of the library was verified using an Agilent Technology 2100 bioanalyzer. Sequencing was performed on an Illumina HiSeq 2500 system at the Cornell University Biotechnology Resource Center (Ithaca, NY, USA)

### Hi-C library construction and sequencing

For Hi-C sequencing, chromatin was isolated, and libraries were prepared from 200 mg of *M. persicae* feeding on *N. tabacum* using the animal Proximo Hi-C kit (Phase Genomics, https://phasegenomics.com/) according to the manufacturer’s instructions. Briefly, insect tissues were ground in liquid nitrogen and crosslinked in crosslinking solution, at 23°C for 20 min. The crosslinking reaction was stopped in 100 µl of quenching solution at 23°C for 20 min followed by centrifugation at 17,000 × g for 5 min. Insect cells were lysed in 700 µl of animal lysis buffer 1, washed in 500 µl of Tris-buffered saline (TBS, 50 mM Tris-HCl, 150 mM NaCl, pH 7.5) buffer, followed by incubating in 500 µl of animal lysis buffer 2 and another washing step by 500 µl of TBS buffer. The chromatin was then incubated at 37°C with 300 µl of proximo fragmentation buffer and 17 µl of proximo fragmentation enzyme for 1 hour, followed by centrifugation at 17,000 × g for 5 min. The pallet was dissolved in 500 µl of deionized water.

One µg of DNA was then incubated at 23°C with 1500 µl of proximity-ligation buffer and 10 µl of proximity-ligation enzyme in a final volume of 2 mL for 4 h. After reverse crosslinking, DNA was column-purified, Hi-C junctions were bound to streptavidin beads, and beads were washed to remove unbound DNA. Washed beads were then used to prepare paired-end deep sequencing libraries using proximity library preparation reagents. The proximity Hi-C libraries were then subjected to PCR amplification, followed by DNA purification using solid phase reversible immobilization (SPRI) beads (Beckman Coulter Life Sciences, https://www.beckman.com/)and resuspended in 30 µl of elution buffer. The Hi-C libraries were sequenced at the Biotechnology Resource Center at Cornell University (Ithaca, NY, USA) using the NextSeq500 platform (Illumina) to obtain 76 bp paired-end reads.

### Transcriptome sequencing

To aid in gene prediction, the *M. persicae* transcriptome was sequenced by Illumina strand-specific RNA-Seq and PacBio Iso-Seq platforms. Total RNA was extracted from mixed-instar aphids feeding on *N. tabacum* using the SV Total RNA isolation kit (Promega, Madison, WI, USA). Briefly, 100–120 mg of insect tissue was grinded in liquid nitrogen using a mortar and pestle and incubated at 70°C in RNA lysis buffer (4 M guanidine thiocyanate; 0.01 M Tris, pH 7.5; 0.97% β-mercaptoethanol [added right before use]) for 3 min. This solution was centrifuged for 10 min at 14,000 × g. The supernatant was mixed with RNA wash solution (60 mM potassium acetate; 10 mM Tris-HCl, pH 7.5; 60% ethanol) and passed through a spin column provided with the kit. To remove DNA, DNase incubation mixture (40 μl of yellow core buffer, 5 μl of 0.09 M MnCl_2_, 5 μl of DNase I) was added to the column membrane and incubated at 23°C for 15 min. The reaction was stopped by adding 200 μl DNase stop solution to the column membrane, followed by washing twice with RNA wash solution, and RNA was dissolved in 50 μl of nuclease-free buffer. Strand-specific RNA-Seq libraries were constructed using a previously described protocol (Zhong et al., 2011) and sequenced at the Cornell University Biotechnology Resource Center (Ithaca, NY, USA) on an Illumina HiSeq 2500 sequencing system. For Iso-Seq, 20 μg of RNA isolated from mixed-instar aphids using SV total RNA isolation kit (Promega) as described above, was proceed at Duke Center for Genomic and Computational Biology (Durham, NC, USA) for PacBio large-insert library construction and sequencing using standard SMRTbell template preparation kits. The library insert size ranged from 500 to 4,500 bp and one SMRT cell was run on the PacBio Sequel platform.

### Genome assembly

The PacBio long reads were corrected and assembled by Falcon (Chin et al., 2016). The resulting contigs were first polished by aligning the raw PacBio reads to the assembly and correcting the sequencing errors using Arrow (Pacific Biosciences). To further improve the assembly, another round of polishing was performed by aligning the Illumina reads to the contigs and correcting the errors using Pilon (Walker et al., 2014). The contigs were then blast-compared against the NCBI non-redundant nucleotide database using BLASTN to identify and remove contamination. To remove the redundant sequences caused by the heterozygosity, the contigs were blast-compared against each other. For each individual contig, if 90% of the length was covered by another, longer contig, then this short contig was considered as redundant, and removed. Pseudo-chromosomes were constructed with Hi-C data using the 3D-DNA pipeline (Dudchenko et al., 2017). The Hi-C reads were aligned to the polished contigs using the Juicer pipeline (Durand et al., 2016). The 3D-DNA pipeline was run with the following parameters: -i 1 -r 5.

### Annotation of repetitive elements

MITE (miniature inverted-repeat transposable elements) were identified from the genome assembly using MITE-Hunter (Han and Wessler, 2010). We generated a *de novo* repeat library by scanning the assembled genome using RepeatModeler (http://www.repeatmasker.org/RepeatModeler), and classified repeats with the RepBase library (Jurka et al., 2005). We subsequently compared these repeat sequences against the NCBI non-redundant (nr) protein database using BLASTX with an e-value cutoff of 1e-5, and those having hits to known protein sequences were excluded. Finally, we identified repeat sequences by scanning the assembly using the *de novo* repeat library with RepeatMasker (http://www.repeatmasker.org/) and the RepeatRunner subroutine (http://www.yandell-lab.org/software/repeatrunner.html) in the MAKER annotation pipeline (Cantarel et al., 2008)

### Gene prediction and annotation

Protein-coding genes were predicted from the genome assembly using the automated pipeline MAKER (Cantarel et al., 2008). The evidence that was used included complete aphid coding sequences collected from NCBI, transcripts assembled from our strand-specific RNA-Seq data, high quality transcript sequences from Iso-Seq, completed proteomes of *Acyrthosiphon pisum, Aphis glycines, Diuraphis noxia, Myzus cerasi, Myzus persicae*, and *Rhopalosiphum padi*, and proteins from the Swiss-Prot database. All of these sequences were aligned to the genome assembly using Spaln (Gotoh, 2008). MAKER was used to run a battery of trained gene predictors, including Augustus (Stanke and Waack, 2003), BRAKER2 (Bruna et al., 2021), and GeneMark-ET (Lomsadze et al., 2014), and then integrated the experimental gene evidence to produce evidence-based predictions. To functionally annotate the predicted genes, their protein sequences were compared against different protein databases including UnitProt (TrEMBL and SwissProt) and two insect proteomes (pea aphid and psyllid) using BLAST with an e-value cutoff of 1e-4.

### Identification of horizontally transferred genes

To identify HTGs in *M. persicae* strain USDA-Red (Ramsey et al., 2007; Ramsey et al., 2014) we compared this lineage to six databases of complete proteomes in UniProt, including archaea, bacteria, fungi, plants, metazoa (excluding proteins from species in the Arthropoda), and other eukaryotes (the remaining eukaryotes excluding fungi, plants, and metazoa). The index of horizontal gene transfer, h, was calculated by subtracting the bitscore of the best metazoan match from that of the best non-metazoan match as described by Crisp et al. (2015). We defined candidate HTGs as those with h ≥ 30 and the bitscore of the best non-metazoan match hit ≥ 100. The corresponding scaffold sequences of these candidates were BLAST-compared against NCBI nt database to exclude contamination. We then performed phylogenetic analyses to validate HTGs. For each candidate gene, its protein sequence was compared by BLASTP against the protein databases of six taxa (archaea, bacteria, fungi, plants, metazoan, and other eukaryotes).

The top five hits from each taxon were extracted, and aligned with the protein sequence of the candidate gene using MUSCLE (Larkin et al., 2007). Each alignment was trimmed to exclude regions where gaps were more than 20% of sequences. Phylogenetic trees were constructed with PhyML (Guindon et al., 2009) using a JTT model with 100 bootstraps. HTGs were considered validated if the genes were monophyletic with taxa of bacteria, fungi, or other microorganisms.

To identify HTGs in *M. persicae* strain G006 (Ramsey et al., 2007; Mathers et al., 2017), we implemented *de novo* HTG annotation following the same protocol as for *M. persicae* strain USDA-Red. The identification of whitefly HTGs has been described previously (Chen et al., 2016). We conducted a homology search using the HTGs that we found in *M. persicae* strain USDA-Red and other genes previously reported in aphids (Parker and Brisson, 2019; Verster et al., 2019). The HTG homologs in other aphid species (*A. pisum, A. glycines, Aphis gossypii, Daktulosphaira vitifoliae, D. noxia, E. lanigerum, M. cerasi, Pentalonia nigronervosa, Rhopalosiphum maidis*, and *R. padi*) were identified using reciprocal homology blast (Table 1; gene IDs in other aphid species are listed in Table S1). The newest-version genomes of all aphid species annotated were downloaded from AphidBase (https://bipaa.genouest.org/is/aphidbase/).

### Design of dsRNA and cloning of target HTG sequences in the TRV-VIGS vector

For each target gene, dsRNA fragments with a length of 150 to 300 bp were designed using the ERNAi website (https://www.dkfz.de/signaling/e-rnai3/) (Horn and Boutros, 2010). To reduce the chances of off-target silencing, the designed fragments were checked against the reference genomes of the experimental species, *M. persicae* strain USDA-Red (this study), *B. tabaci* MEAM1 (Chen et al., 2016), *N. benthamiana* (Bombarely et al., 2012), and *C. septempunctata*, using a VIGS tool (Fernandez-Pozo et al., 2015) and local BLAST (Mount, 2007) to eliminate dsRNA fragments that contain any full matches of ≥ 19 nt for bacterial and plant origin HTGs and > 21 nt for fungal origin HTGs to the reference genomes. The retained dsRNA fragments were cloned into the TRV-VIGS vector (Senthil-Kumar and Mysore, 2014) using the Invitrogen Gateway recombination cloning system (Invitrogen, USA). Briefly, dsRNA fragments were first PCR-amplified from aphid cDNA using primers designed based on the dsRNA fragments (Table S2). PCR products were then cloned into pDONR207 using the Gateway BP clonase, dsRNA fragments in pDONR207 were swapped into Gateway compatible TRV2 plasmid (pTRV2) by recombination using Gateway LR clonase. The final constructs were named TRV2-GOI (gene of interest). In parallel, we obtained a construct TRV2-PDS, which carries a dsRNA fragment targeting *N. benthamiana* phytoene desaturase as a positive control for expression silencing (Velasquez et al., 2009). As negative controls, we used TRV2-GFP or TRV2-GUS, which carry dsRNA fragments targeting Green Fluorescent Protein (GFP) or *Escherichia coli* beta-glucuronidase (GUS) genes, respectively, but not sequences that are found in either the target insects or *N. benthamiana* (Vaghchhipawala et al., 2011), as well as a TRV2-EV (empty vector).

### *Transient* Agrobacterium *infection of* N. benthamiana *for TRV-based VIGS*

To infiltrate plants, plasmids (TRV1 and TRV2 carrying the gene of interest) were transformed into *Agrobacterium tumefaciens* and then infiltrated into *N. benthamiana* as previously described (Senthil-Kumar and Mysore, 2014). Briefly, the TRV1 and TRV2-GOI plasmids were transformed into *A. tumefaciens* strain GV3101. Each TRV2-GOI *Agrobacterium* culture was mixed with one carrying TRV1 and adjusted to a final OD_600_ of 0.3 with cell suspension buffer (final concentrations at 0.01 M MES, 0.01 M MgCl_2_, and 0.2 mM acetosyringone). After three hours of incubation at 23°C, the *Agrobacterium* TRV mixtures were infiltrated to saturate three leaves of v4 stage *N. benthamiana* using 1-ml needleless syringes. Inoculated plants were kept in a growth chamber for 2-3 weeks, at which point the TRV-PDS infiltrated plants showed photobleaching symptoms that indicated viral spread. The TRV-GOI plant leaves were assayed for the presence of TRV1 and the expression of GOI dsRNA using PCR (Figure S1). Leaf samples were collected for RNA extraction using the SV Total RNA Isolation system (Promega, USA) and cDNA synthesis using the High-Capacity cDNA Reverse Transcription Kits (Applied biosystems, USA). The synthesized cDNA samples were amplified by PCR using Phusion™ High-Fidelity DNA Polymerase (ThermoFisher Scientific, USA) and the VIGS primers (Table S2, Figure S2, S3). A 20 µl PCR reaction contains 4 µl betaine, 4 µl 5 x High-Fidelity buffer, 1 µl 2.5 mM dNTPs, 0.2 µl Phusion polymerase, 1 µl each of 10 mM forward and reverse primers, 1 µl cDNA and 7.8 µl of ddH_2_O. The PCR program was 95°C for 3 mins, followed by 25 cycles of 95°C for 30 s, 60°C for 30 s, and 72°C for 1 min, with a final step of 72°C for 5 min. The confirmed TRV-GOI plants were then used for aphid bioassays.

### Aphid bioassays

For each GOI, 5 or 6 infiltrated plants were used for aphid bioassays, and two aphid cages were attached to the adaxial side of fully developed young leaves on each plant. To set up the bioassays, ten adult aphids were added to each cage and allowed to produce nymphs for 24 h. The adults were removed, and 25 newborn nymphs were scored for survival every 24 hours for up to five days. After five days, 10-15 adult aphids were collected and flash frozen in liquid nitrogen for target gene expression analyses using quantitative reverse transcriptase-PCR (qRT-PCR) (Primers see Table S2, Figures S2, S3).

To test the effects of silencing bacterial and plant-origin HTGs on aphid reproduction, immediately after the 5^th^-day survival check, the cages were moved to young leaves and five adult aphids were left in each cage to measure aphid reproduction for a week. To test the effects of dsTor and dsCarRP on aphid reproduction, one adult aphid was added to each cage for 24 h, and then the adult aphid was removed. Nymphs that were deposited by the adult aphids were collected after 7 days, the number of progeny and the survival of the nymphs was scored, and adults were collected for qRT-PCR analysis.

### Ladybug bioassays

On wildtype *N. benthamiana* plants, *M. persicae* prefer feeding on older, senescing leaves but the TRV virus tends to move to younger and new leaves. To generate enough aphids for the ladybug bioassays, we performed additional VIGS experiments and maintained a larger aphid population on *asat2-1* mutant *N. benthamiana* plants (Feng et al., 2021), which are deficient in acylsugars and allow *M. persicae* feeding on the TRV-infected younger leaves.

For ladybug bioassays, two-day-old *C. septempunctata* larvae were used for all experiments. Each ladybug larva was individually caged in vented plastic cups and aphids from VIGS *N. benthamiana* plants were served every other day. The survival of each ladybug was tracked for seven days, and ten ladybugs were monitored for each of the VIGS treatments. The experiments were repeated twice using wildtype and twice with *asat2-1* mutant *N. benthamiana*. To confirm tri-trophic persistence of RNAi signals, we also conducted assays using *C. montrouzieri* adults.

### Whitefly bioassays

As *B. tabaci* strain MEAM1 could not survive on wildtype *N. benthamiana*, we used *asat2-1* mutant *N. benthamiana* (Feng et al., 2021) for bioassays. For each experiment, 3 or 4 TRV-GOI infiltrated plants were used for each gene of interest, and three cages were attached to each plant. Ten young adult whiteflies (emerged within the past week) were placed in each cage. After seven days, the survival rates of whiteflies were assessed and compared across different treatments. The surviving whiteflies were collected for q-RT-PCR analyses of target gene expression. The whitefly VIGS bioassays were repeated three times with similar results. Two replicates of a separate experiment were set up to collect whiteflies after one day for gene expression analyses.

### mRNA expression and qPCR analysis

We used q-RT-PCR to measure the expression of target GOIs in treated aphids and whiteflies. Total RNA was extracted from 10-20 *M. persicae* using the SV Total RNA Isolation system (Promega, USA) and RNA was reverse-transcribed using the High-Capacity cDNA Reverse Transcription Kits (Applied biosystems, USA). cDNAs were then diluted 10-fold and used for qPCR reactions. Each qPCR reaction contained 5 μl of the PowerUp™ SYBR™ Green PCR master mix (Applied Biosystems, USA), with 1 μl of each qPCR primer (Table S2) and 1 μl of cDNA. qPCR primers were designed to not overlap with the selected dsRNA in each of the targeted genes to avoid detecting signals from ingested dsRNAs. PCR reactions were initiated with an incubation at 95°C for 30 seconds, followed by a 40 times cycle of 95°C for 5 seconds, 60°C for 1 minute, and a melting curve was collected after the reaction. The Ct values were used to quantify and analyze gene expression according to the 2^-^ΔΔCT method (Livak and Schmittgen, 2001). Ubiquitin and/or *EF1a* were used as the internal control genes in each qPCR experiment for bacterial and plant origin HTGs and *RpL7* was used as an internal control for *Tor* and *CarRP*. Changes in the expression levels of each gene were calculated by comparing the ratio of the ΔΔCT values of samples from aphids and whiteflies fed on TRV2-GOI plants to those fed on TRV2 empty vector control plants.

### Data analyses and statistics

For both insect bioassays and gene expression assays, data were pooled from multiple experiments for statistical analyses. We tested for differences using the Univariate analysis of variance with a fixed factor of treatments and a random factor of experiment, followed by a Bonferroni post-hoc test for multiple comparisons using SPSS v.25 (IBM, Armonk, NY). Differences in ladybug survival were tested using the Cox mixed-effect model (Therneau and Grambsch, 2000) followed by a Tukey post hoc test using R Studio v 1.3.959 (RStudio Team, 2020). For dsTor and dsCarRP, the aphid reproduction data were tested using ANOVA followed by the Tukey’s HSD test. For gene expression data, Mann-Whitney U-tests were used to test for differences.

## Results

### *Chromosome-scale assembly of the* M. persicae *genome*

The assembled *M. persicae* genome has a total length of 383.0 Mb and consists of 331 contigs with N50 length of 4.5 Mb. A total of 364.4 Mb (95.1% of the assembly) are clustered into six chromosomes (Figure 1), which is consistent with the commonly observed 2n = 12 karyotype of *M. persicae* (Blackman, 1980). To evaluate the completeness of the *M. persicae* genome assembly, the Illumina sequencing reads were aligned to the assembly, allowing up to three mismatches using BWA-MEM (Li and Durbin, 2009). Of the Illumina reads, 95.2% could be mapped to the assembly, indicating that most of the acquired reads were successfully assembled into the genome. Furthermore, the completeness of the genome assembly, as evaluated by benchmarking universal single-copy orthologs using BUSCO v 3.0.2 (Simao et al., 2015), showed that 94.2% of the core eukaryotic genes were at least partially captured by the genome assembly and 92.6% were completely captured. We predicted a total of 27,430 protein-coding genes in the *M. persicae* genome. The assembled genome has been deposited in GenBank under PRJNA888091.

**Figure 1.**
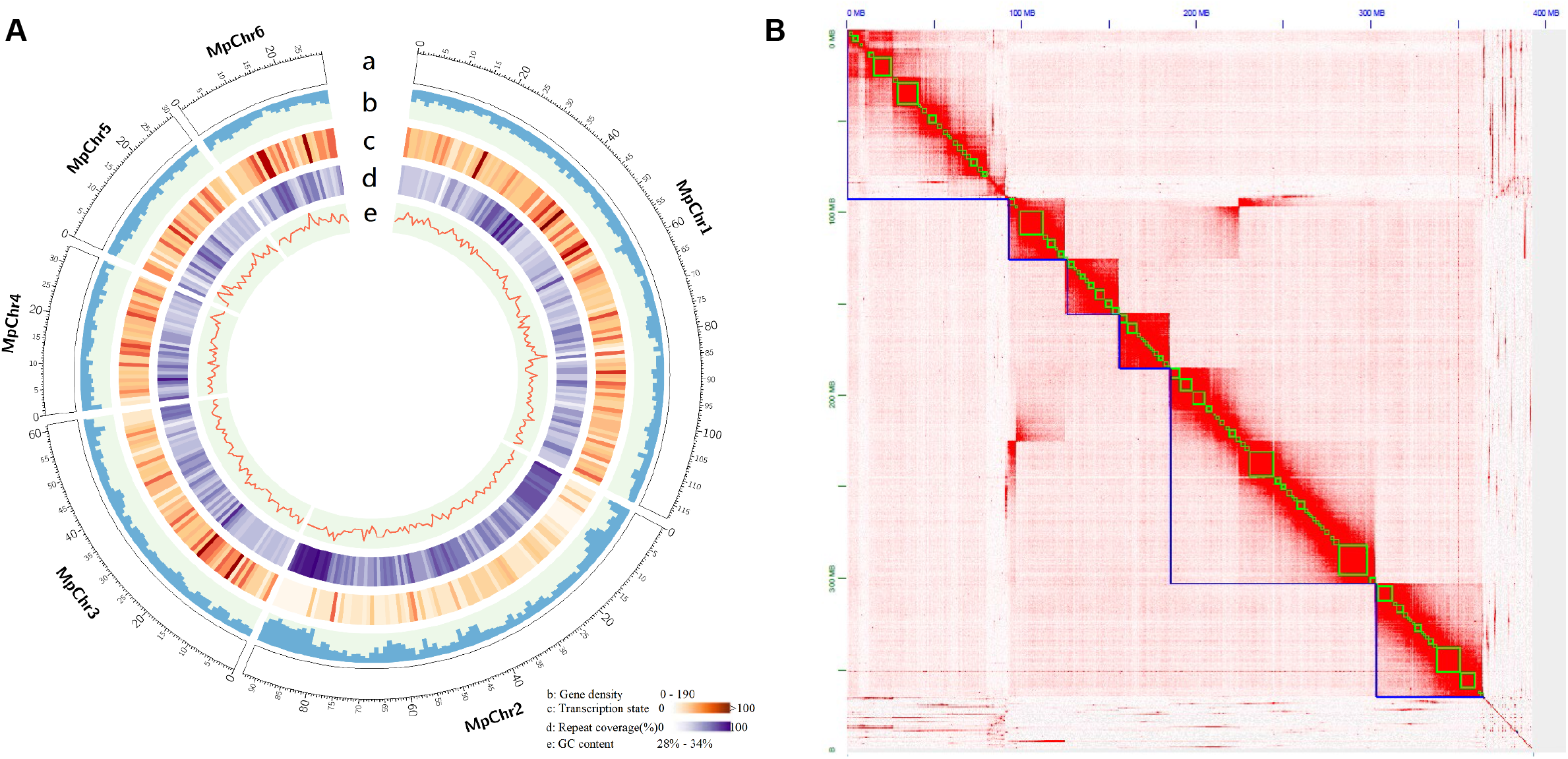
Chromosome-scale genome assembly of a tobacco-adapted red *Myzus persicae* strain. (**A**) *M. persicae* genome landscape. (a) Ideogram of the six *M. persicae* pseudochromosomes at the Mb scale. (b) Gene density is represented as number of genes per Mb. (c) Transcription state. The transcription level was estimated by read counts per million mapped reads in 1-Mb windows. (d) Percentage of coverage of repeat sequences per Mb. (e) Guanine-cytosine (GC) content in 1-Mb windows. (**B**) Heatmap showing frequency of HiC contacts along the *M. persicae* genome assembly. Blue lines indicate super scaffolds and green lines show contigs.

### *Horizontally transferred genes (HTGs) in* M. persicae *str. USDA-Red*

We identified 30 genes that were predicted to be horizontally transferred from bacteria, viruses, fungi, or plants into *M. persicae* strains USDA-Red and G006 (Table 1 and Table S1). Genes of bacterial origin include an uncharacterized protein with predicted ATP binding function, N-acetylmuramoyl-L-alanine amidase (amiD), and a microcin c7 self-immunity protein, which is a homolog of the *A. pisum* murein tetrapeptide carboxypeptidase (ldcA1, ACYPI009109) (Table 1). Genes of viral origin were all identified based on homology search using previously reported HTGs (Parker and Brisson, 2019; Verster et al., 2019). For the non-structural protein NS1 superfamily genes, three copies were identified in our *M. persicae* USDA-Red genome, whereas either one or two copies were present in other species, including the *A. pisum* genome where these HTGs were initially identified (Parker and Brisson, 2019). Fungal origin genes were mainly predicted to be involved in carotenoid biosynthesis, with nine copies of lycopene cyclase-phytoene synthase and four copies of carotenoid desaturase (Table 1). Genes of plant origin are most similar to genes from microalgae and encode uncharacterized proteins with predicted ankyrin repeats, i.e. MPE002740, MPE008575, MPE008576, MPE010373, MPE013670, MPE016359, MPE018635, MPE018636, MPE023973 (Table 1). In addition, one other plant-origin HTG, MPE011636, was identified from our *de novo* annotation (Table 1).

Genomic comparisons across different aphid species (*A. pisum, A. glycines, A. gossypii, D. vitifoliae, D. noxia, E. lanigerum, M. cerasi, P. nigronervosa, R. maidis*, and *R. padi*) showed that most HTGs from *M. persicae* strains USDA-Red and G006 are present in multiple other aphid species (Table 1). For instance, the fungal-origin lycopene cyclase-phytoene synthase and all of its duplicates were found in all assessed aphid species. By contrast, one of the plant-origin uncharacterized ankyrin repeat proteins (MPE0018635) and the two additional copies of non-structural protein (MPE014463 and MPE006690) were found only in *M. persicae* strains G006 and USDA-Red. (Table 1).

### Tri-trophic persistence of RNA

To examine the possibility of tri-trophic persistence of RNAi signals, we used two ladybug species, *C. septempunctata* larvae (Figure 2) and *C. montrouzieri* adults (Figure S4). A dsRNA fragment (dsGFP), targeting the green fluorescent protein gene which it is not naturally present in plants, aphids, or ladybugs, was expressed in *N. benthamiana*. The plant-expressed dsRNA of GFP fragments were consistently detected in both ladybug species that were fed on aphids that fed on dsGFP plants (Figure 2 and Figure S4). As the aphids were removed from the plants for feeding *C. septempunctata* larvae and *C. montrouzieri*, we could rule out direct transfer of the RNA from plants to beetles. Given the observed transfer of RNA from plants to beetles via aphids, we proposed using RNAi to target horizontally transferred genes that are specific to insect pests, thereby avoiding negative impacts on non-target species, in particular beneficial insects like *C. septempunctata* larvae and *C. montrouzieri*.

**Figure 2.**
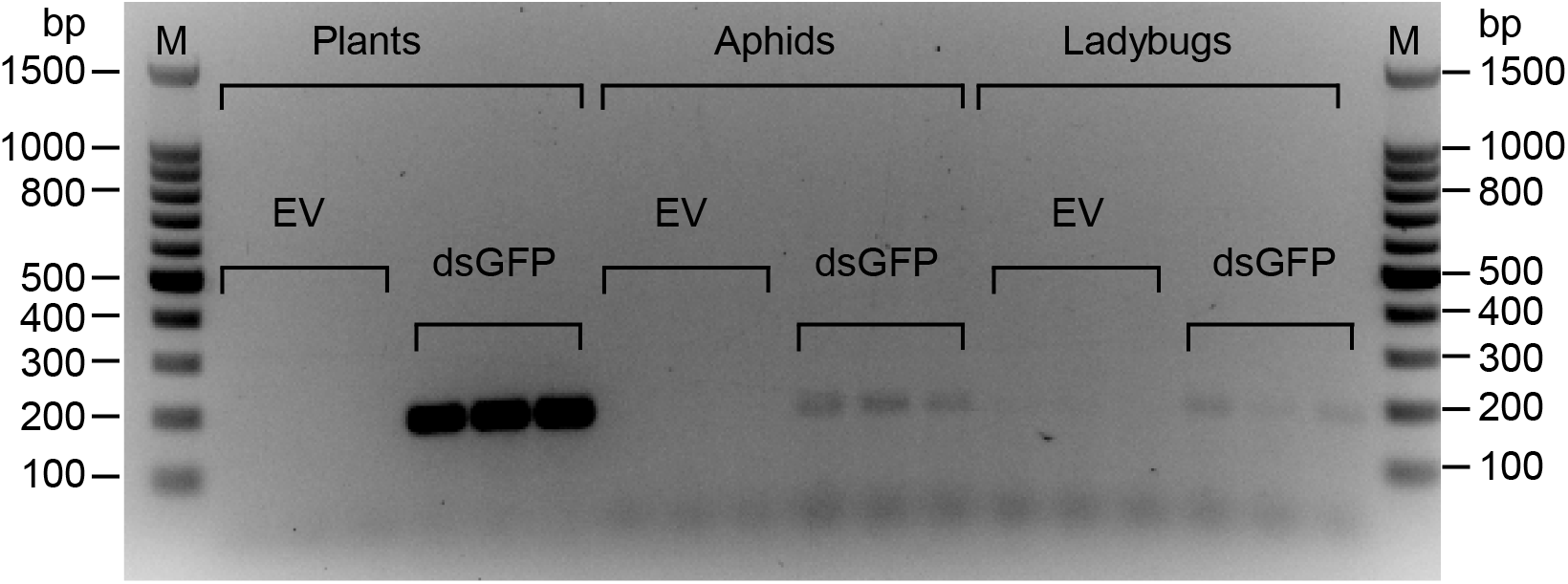
Evidence of tri-trophic persistence of dsRNA. Agarose gel image showing the detection of dsGFP in *Coccinella septempunctata* ladybug larvae after consuming *Myzus persicae* aphids that had fed on *Nicotiana benthamiana* infiltrated with TRV2-*GFP*. The expected amplicon size is 180 bp, which is consistent to the band size shown in gel. PCR from cDNA samples of the following treatments shown in each lane: Plants: plants that infiltrated with TRV2 empty vector (EV) or TRV2-*GFP* vector (dsGFP); Aphids: aphids fed on EV and dsGFP VIGS plants; Ladybugs: ladybugs fed on aphids that were fed on EV and dsGFP VIGS plants. Each treatment was performed with three biological replicates (as shown as three lanes), and this experiment was repeated twice. M: Promega 100 bp DNA ladder.

### *Silencing of* M. persicae *bacterial origin HTGs using VIGS*

To evaluate the effects of silencing bacterial-origin HTGs (Amidase, ATP binding protein, and *ldcA*) on aphids and the ladybugs consuming them, we knocked down gene expression in *M. persicae* using TRV VIGS. Expression of all three VIGS-targeted genes was significantly reduced relative to aphids feeding on TRV2-EV, TRV2-*GUS*, and/or TRV2-*GFP* control plants (*p* < 0.05) (Figure 3). For the Amidase gene, we observed significantly lower aphid survival as early as 24 hours after the initiation of aphid feeding on VIGS plants, compared to both TRV2-*GUS* and EV controls (*p* < 0.05). In the case of the ATP binding protein, aphid survival decreased significantly on the TRV2-*ATPb* VIGS plant compared to TRV2-*GFP* (only initially at 24 hours post feeding) or EV plants (throughout the time period monitored, from 24 hours to 120 hours post-feeding) (*p* < 0.05). In the case of *ldcA*, we observed significantly lower aphid survival on TRV2-*ldcA* plants comped to both TRV2-*GFP* and EV controls (*p* < 0.05). We did not observe significant effects on aphid reproduction with any of the three tested genes. Furthermore, there were no significant effects on the survival of *C. septempunctata*, when using either wildtype or *asat2-1* mutant *N. benthamiana* plants for the VIGS experiments.

**Figure 3.**
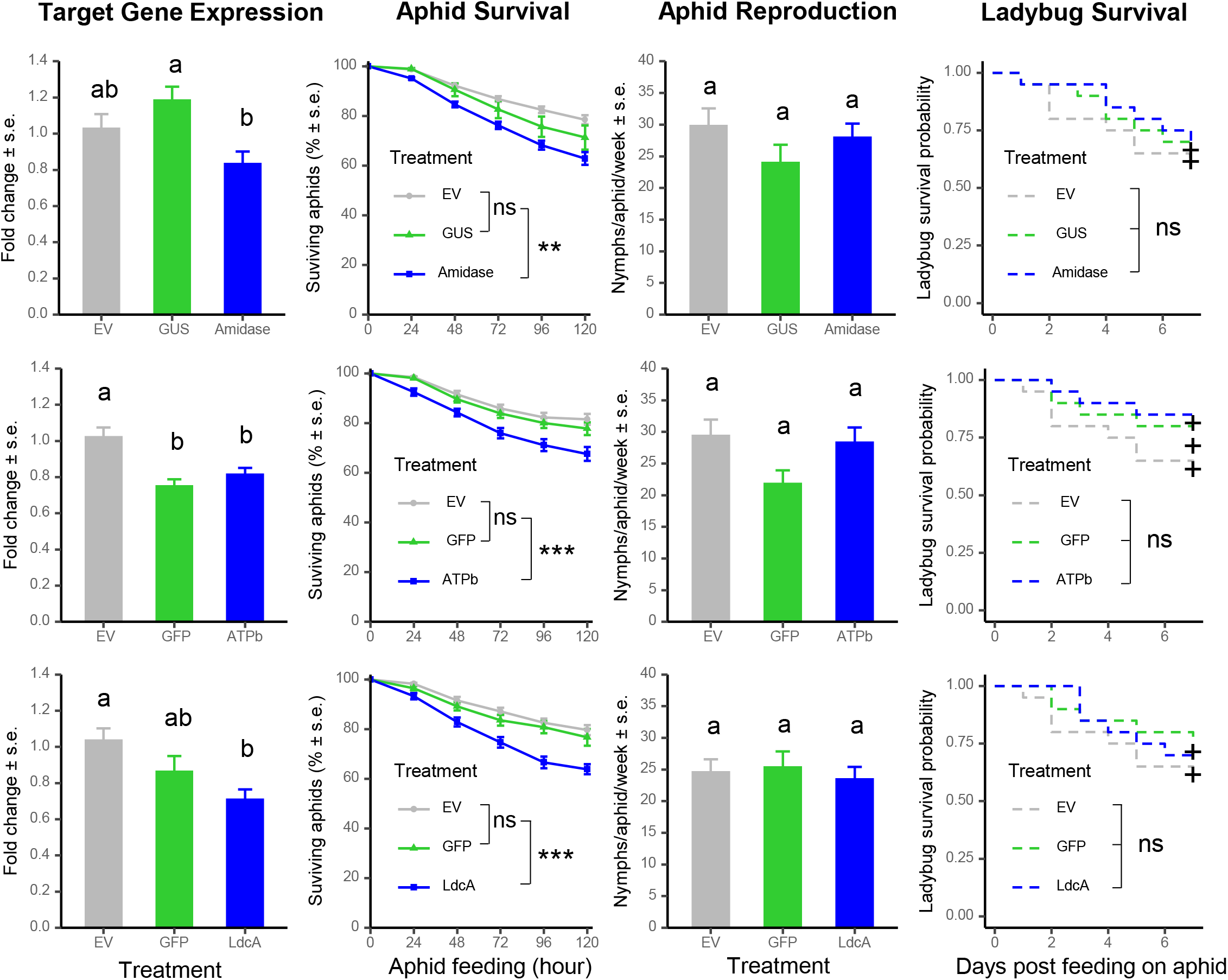
Knock-down of bacterial-origin horizontally transferred genes reduced aphid survival with no effects on ladybug survival. Bioassay experiments for were replicated at least 5 times, with 5-8 plants for each construct as biological replicates in each experiment. Letters above the bars indicate significant differences between treatments by ANOVA (*p* < 0.05). Significant differences of the aphid survival between treatments were indicated for the last time point (*i*.*e*. 120 hrs), ** p < 0.01, *** p < 0.001, ns: not significant.

### *VIGS-mediated silencing of* M. persicae *plant origin HTGs*

After 120 h of feeding, expression of five plant-origin HTGs, all of which are uncharacterized proteins with ankyrin repeat domains (Table 1), was significantly downregulated by VIGS in comparison to *M. persicae* on TRV2-*GFP* and/or EV control plants (*p* < 0.05) (Figure 4). The effects on gene expression varied from a 20% decrease for MPE018635 to a 50% decrease for MPE013670. We observed significantly lower aphid survival on the VIGS plants compared to both TRV2-*GFP* and EV controls (*p* < 0.05) (Figure 4). The decreases in aphid survival ranged from 30% for MPE018635 to 40% for MPE010373 after 120 hours of aphid feeding (Figure 4). The significant decreases in aphid survival were observed starting from 24 h after initiation of aphid feeding and continue to decrease over time, with the exception of MPE023973, where significant decreases were observed only starting at 96 h (Figure 4). Similar to the bacterial-origin HTGs, we did not observe significant effects on aphid reproduction with any of these five plant-origin HTGs. Similarly, there were no significant effects on the survival of *C. septempunctata*, when using either wildtype or *asat2-1* mutant *N. benthamiana* plants for the VIGS experiments.

**Figure 4.**
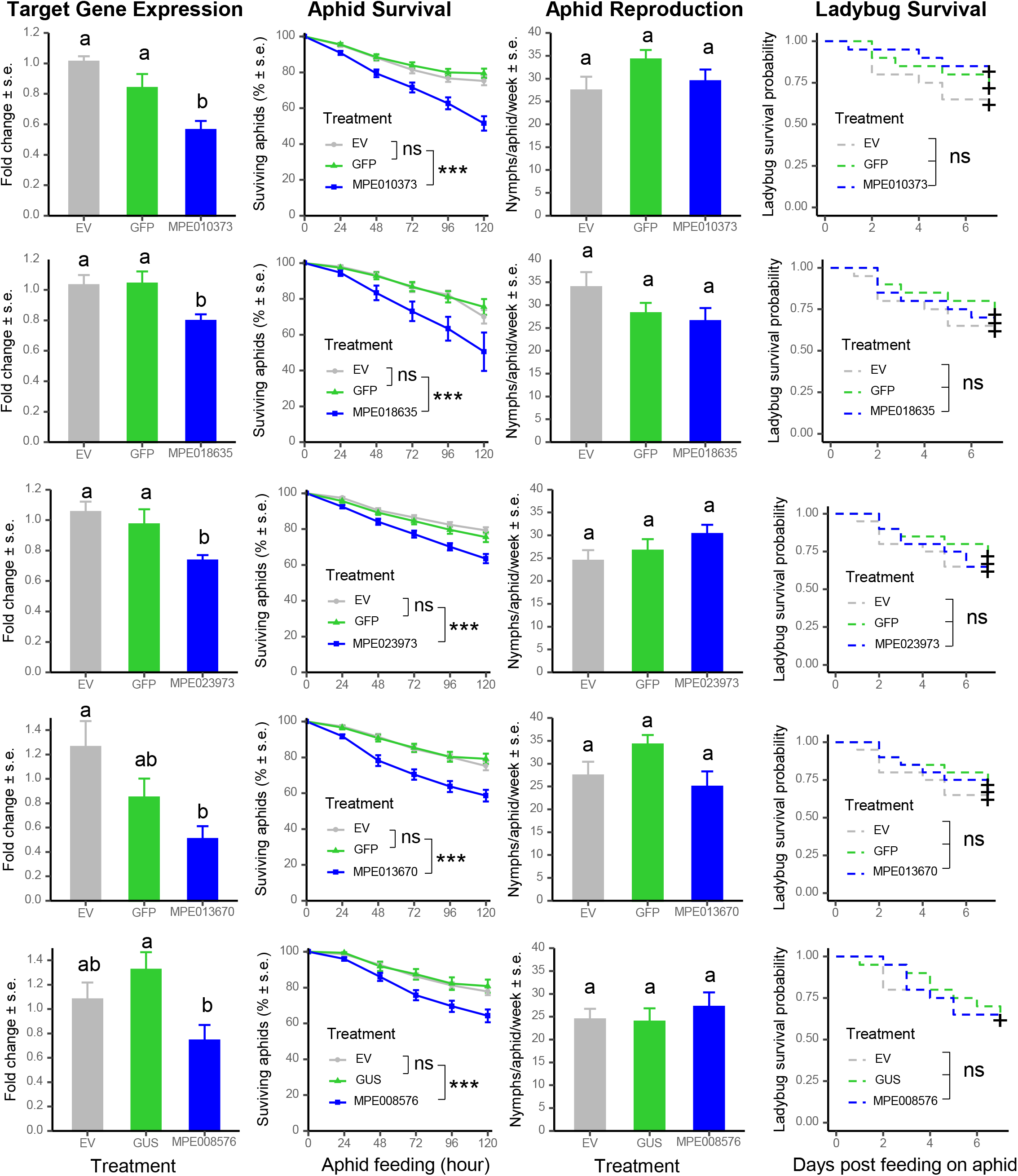
Knock-down plant-origin horizontally transferred genes reduced aphid survival with no effects on ladybug survival. Bioassay experiments for were replicated at least 5 times, with 5-8 plants for each construct as biological replicates in each experiment. Letters above the the bar graphs indicates significant differences between treatments by ANOVA (*p* < 0.05). Significant differences of the aphid survival between treatments were indicated for the last time point (*i*.*e*. 120 hrs), ** p < 0.01, *** p < 0.001, ns: not significant.

For three additional genes of plant origin, MPE018636, MPE002740, and MPE016359, we didn’t observe significant effects on aphid survival. Among five VIGS experiments, we only observed significant aphid survival decrease on dsMPE018636 VIGS plant in one of the experiments, an experiment in which we did not observe a significant decrease in expression of the targeted gene.

### VIGS-mediated silencing of the fungal origin Tor and CarRP genes

Carotene desaturase (*Tor*) is required for the production of the red carotenoid, torulene, and carotenoid cyclase–carotenoid synthase (*CarRP*) catalyzes both the committed step for carotenoid biosynthesis and later steps involving the formation of cyclic carotenoids (Hansen and Moran, 2011; Novakova and Moran, 2012). We conducted *Tor* and *CarRP* VIGS experiments using a time point seven days after aphids were born on VIGS plants. Compared to aphids on dsGFP control plants, the expression levels of both genes were reduced by approximately 50% (*p* < 0.05) in aphids feeding on TRV2-*Tor* and/or TRV2-*CarRP N. benthamiana* plants (Figure 5A, B). No apparent macroscopic phenotypic changes (particularly loss of red color) were observed among the groups of aphids.

**Figure 5.**
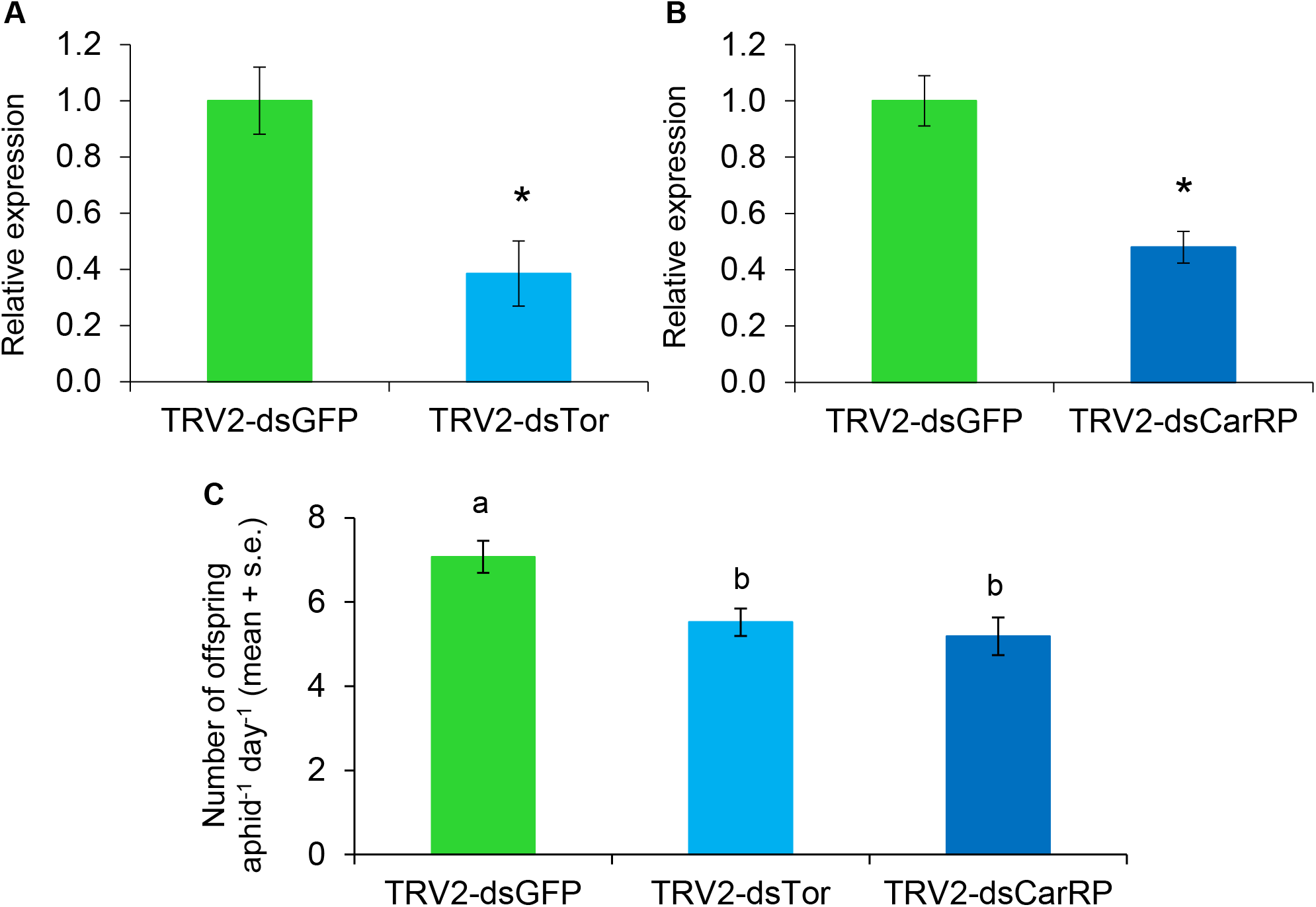
Silencing of *Tor* and *CarRP* genes reduces aphid reproduction. **A**) *Tor* gene and **B**) *CarRP* gene expression after 7 d exposure of *Myzus persicae* to TRV-VIGS in *Nicotiana benthamiana*. Data shown are the mean +/- standard error of 5 or 6 biological replicates, with three aphids pooled for each replicate. The Y-axis has arbitrary units, with gene expression in the TRV-GFP strain normalized as 1. \**p* < 0.05, Mann-Whitney U-test relative to the control. **C**. Reproductive output per aphid over 8 days of *M. persicae* reared on *N. benthamiana* plants with the two RNAi constructs. ANOVA results are shown, with different letters referring to treatments with significantly different expression by Tukey’s HSD test.

### *VIGS-mediated silencing of* M. persicae *virus origin cdtB*

We also tested one of the viral HTGs, Cytolethal Distending Toxin B (*cdtB*), which was suggested to be involved in aphid resistance to a predatory wasp (Verster et al., 2019). Somewhat unexpectedly, *cdtB* gene expression were consistently observed to be up-regulated in the TRV-*cdtB* plants relative to TRV-*GFP* and EV controls (Figure 6A). This may be indicative of unexplained upregulation of *cdtB* expression as a compensatory response to the attempted RNA interference. We did not observe any effects on aphid survival in our plant-mediated VIGS experiments, which were conducted in the absence of wasp predation (Figure 6B).

**Figure 6.**
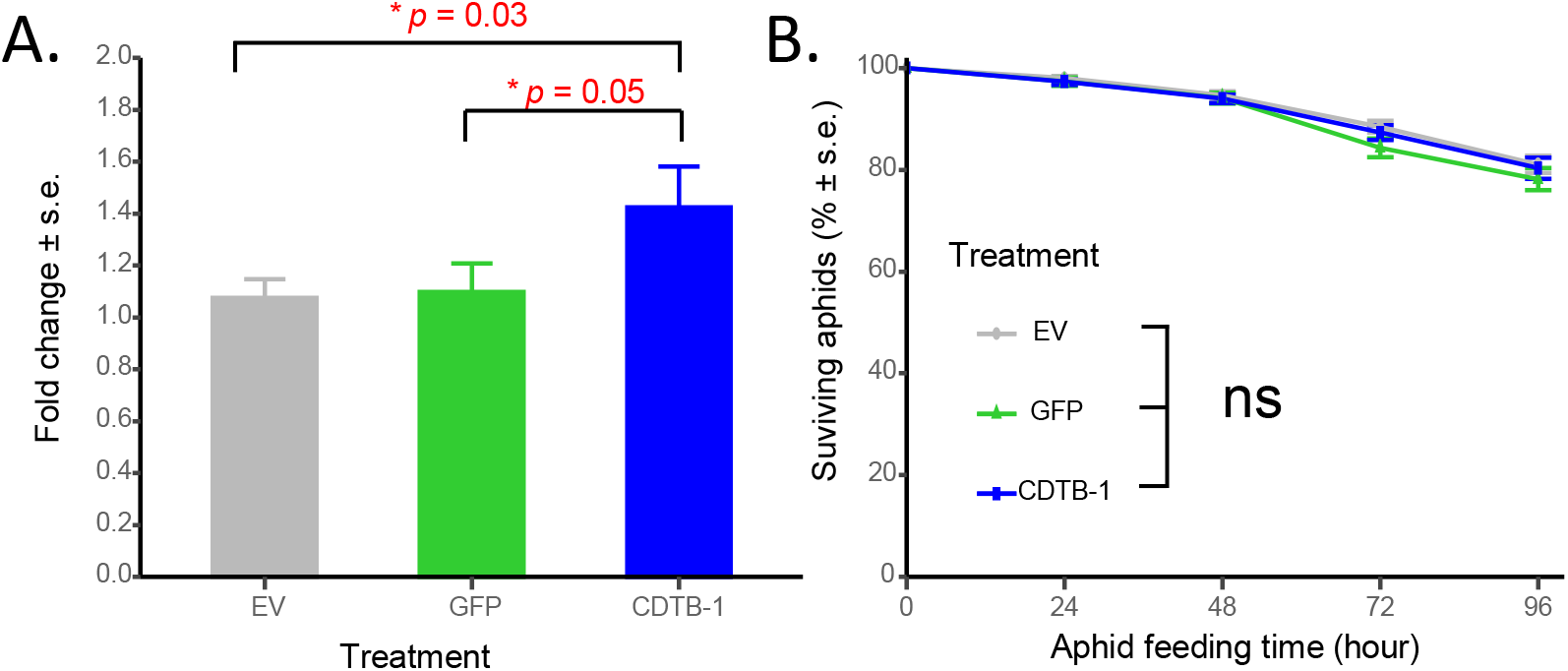
VIGS upregulated *cdtB* (virus-origin HTG) gene expression (A), but did not affect aphid survival (B). Bioassays for were replicated 4 times with similar results. Each experiments included 5-8 plants for each construct as biological replicates. Significant differences between groups were tested using ANOVA with a block effects (experiments), followed by multiple comparisons using Bonferroni method. * p < 0.05, ns: not significant.

### VIGS-mediated silencing of whitefly HTGs

*Bemisia tabaci* MEAM1 has at least 142 HTGs from bacteria and fungi (Chen et al., 2016). As VIGS targets, we chose five *B. tabaci* HTGs, squalene synthase (*SQS, Bta11043*), diaminopimelate decarboxylase (*DAPDC, Bta03593*), argininosuccinate synthase (*ASS, Bta00062*), cyanate hydratase (*CYN, Bta20016*), and aromatic peroxygenase (*APO, Bta04808*). In parallel, we chose three essential whitefly genes as positive controls, including acetylcholine esterase (*AchE*) and toll-like receptor 7 (*TLR7*), which have been targeted previously by RNAi (Malik et al., 2016), and the tubulin-specific chaperon D (*TBCD*), which has been targeted by VIGS in aphids (Guo et al., 2014). We initially conducted VIGS experiments with wildtype *N. benthamiana*, but the low survival of *B. tabaci* on these plants made it impossible to reliably assess the effectiveness gene expression silencing and interpret the results (Figure 7A). Instead, we conducted whitefly VIGS experiments with *asat2-1* mutant *N. benthamiana*, which allow improved whitefly reproduction and survival (Feng et al., 2021). In VIGS experiments with *asat2-1* plants, *AchE* and *SQS* expression was reduced after 1 and 7 days (Figure 7B,C), which resulted in significantly reduced whitefly survival (Figure 7D). Although *TLR7* and *TBCD* gene expression was reduced by VIGS only on day 1 when feeding on *asat2-1* plants (Figure 7B), there was nevertheless a negative effect on whitefly survival over 7 days (Figure 7D). Two possible explanations for this finding are: (i) There was a survivor bias in that we could only measure gene expression levels in surviving whiteflies, and perhaps all whiteflies with efficient *TLR7* and *TBCD* expression silencing were dead after 7 days; or (ii) There may be gene expression compensation at the whole-insect level over time, but not in specific tissues that affect insect survival. Quantitative PCR of fractionated whiteflies would be necessary to determine the time course of *TLR7* and *TBCD* expression silencing and whether VIGS primarily affects gene expression in specific tissue types.

**Figure 7.**
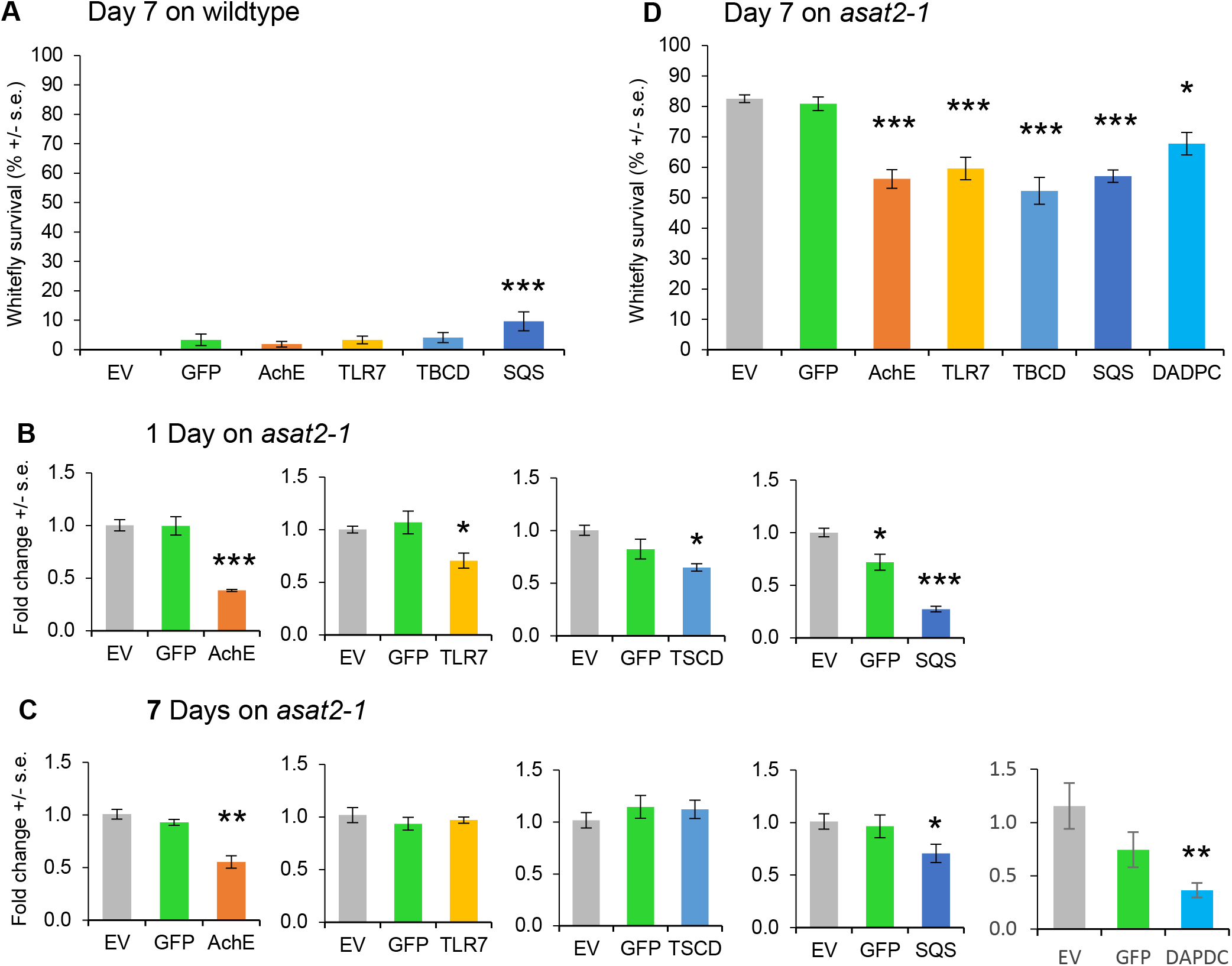
Virus induced gene silencing (VIGS) of whitefly genes using tobacco rattle virus (TRV). *Bemisia tabaci* (whiteflies) were placed on wildtype (**A**) and *asat2-1* (**B, C, D**) *Nicotiana benthamiana* plants infected with TRV expressing VIGS constructs and gene expression was measured after (**B**) one and (**C**) seven days. In each experiment, the empty virus vector (EV) and *GFP*-carrying virus were used as controls. Gene expression was normalized to 1 for EV control plants. Whitefly survival was assessed on TRV-infected plants after seven days of feeding on (**A**) wildtype plants or (**D**) *asat2-1* plants. GFP/AchE/SQS/TLR7/TBCD/DADPC: VIGS plants with TRV expressing RNA constructs targeting GFP, AchE, SQS, TLR7, TBCD, and DADPC respectively; AchE: acetylcholinesterase, SQS: squalene synthase, TLR7: toll-like receptor 7, TBCD: tubulin-specific chaperon D, DADPC: Diaminopimelate decarboxylase. The survival and the 7-day qPCR experiments were repeated three times. The 1-dayqPCR experiments were repeatd twice. Significant differences were determined using one-way ANOVA with a fixed factor of treatments and a block effect of experiment, followed by a Dunnett’s post hoc test for comparing VIGS constructs to the empty vector (EV) control. Mean +/- s.e. of n = 3 (A), n = 9 (B), and n = 27 (C,D). \**p* < 0.05; \*\**p* < 0.01, \*\*\**p* < 0.001.

## Discussion

### Identification of HTGs in insect pests

With the increasing number of available genomes, HTGs have been identified in all branches of life, including a broad range of agricultural insect pests such as aphids and whiteflies (Soucy et al., 2015; Husnik and McCutcheon, 2018). Many of those HTGs have been shown to be expressed and functional after the incorporation into the recipient genome (Wybouw et al., 2016; Husnik and McCutcheon, 2018), providing the recipient species benefits such as nutrition, protection and adaptation to environmental stresses (Soucy et al., 2015; Husnik and McCutcheon, 2018). HTGs can help the recipient insects to digest plant cell wall components that would otherwise be non-digestible (Pauchet and Heckel, 2013; Kirsch et al., 2014; Pauchet et al., 2014; Zhao et al., 2014). For example, the pectin-degrading polygalacturonases found in herbivorous beetles are ancient HTGs from fungi that help beetles to digest cell wall components such as pectin, cellulose, and hemicellulose (Kirsch et al., 2014). HTGs can also help recipient insects detoxify plant defensive metabolites (Daimon et al., 2008). For instance, mulberry latex, such as 1,4-dideoxy-1,4-imino-D-arabinitol (D-AB1) and 1-deoxynojirimycin (DNJ) are highly toxic to non-mulberry specialist insects, but present no harm to the mulberry specialist silkworm, *Bombyx mori*. A beta-fructofuranosidase gene found horizontally transferred from bacteria is highly expressed in the *B. mori* gut and help digesting plant-defensives alkaloidal sugars (Daimon et al., 2008). In sap-feeding insects, HTGs from bacteria have been found to complement the function of their obligate endosymbiont bacteria in the biosynthesis of essential nutrients, such as essential amino acids and vitamins (Husnik et al., 2013; Sloan et al., 2014; Luan et al., 2015). Some HTGs have been proposed to be involved in protecting recipient insects from pathogen or predators (Verster et al., 2019). For example, the *cdtB* gene of bacteriophage origin in the vinegar flies and aphids may function in defending these insects against natural enemies (Verster et al., 2019). As these nutritional and protective functions shown above are critical to insect development, survival, and reproduction, and most of them are species- or genus-specific, those HTGs in insect pests should be further explored as efficient and specific targets for the control of pest species.

### HTGs in aphids and their potential for aphid control

As the genome resources becoming available (International Aphid Genomics Consortium, 2010), many HTGs have been described in aphid genomes (Nikoh and Nakabachi, 2009; Moran and Jarvik, 2010; Nikoh et al., 2010; Parker and Brisson, 2019; Verster et al., 2019; Ding et al., 2020).

Aphids acquired carotenoid biosynthesis genes through HTG from fungi. Carotenoids determine aphid body color, which further influences the ecological interactions between aphids and their predators, such as wasps (Moran and Jarvik, 2010). Consistent to these previous experiments, our plant-mediated VIGS experiment showed negative effects on aphid performance, in this case progeny production. As Tor and CarRP are believed to be dispensable during aphid development, there may be no selection against aphids with reduced expression levels. The absence of visible changes in aphid body color may indicate that expression of the targeted *Tor* and *CarRP* genes is not limiting for aphid carotenoid production, or that the reduced expression resulting from VIGS is not enough to result in phenotypic changes.

In addition to HTGs from fungi, which confer aphid phenotypes that are ecologically relevant, we used plant-mediated VIGS to determine that three HTGs from bacteria influence *M. persicae* survival. Transiently knocking down *ATPb, amiD*, and *LdcA* gene expression significantly reduce aphid survival (Figure 2). This result for *amiD* and *LdcA* in *M. persicae* is consistent with an investigation of these two genes in pea aphids, where knocking down *amiD* and *ldcA* by dsRNAs in artificial diet reduced aphid growth (Chung et al., 2018).

All of the plant-derived HTGs that we identified in *M. persicae* encode uncharacterized proteins with ankyrin repeat domains. The ankyrin repeat domains consists of ∼33 amino acid tandem motifs, which have been well documented in protein-protein interaction studies (Sedgwick and Smerdon, 1999). Since the first report of ankyrin repeat proteins in yeast (*Saccharomyces cerevisiae*), fruit flies (*Drosophila melanogaster*), and nematodes (*Caenorhabditis elegans*) (Breeden and Nasmyth, 1987), ankyrin repeat proteins have been identified in numerous organisms, ranging from viruses and bacteria to humans (Sedgwick and Smerdon, 1999). Bacterial ankyrin repeat proteins are mainly found in species that are obligately or facultatively associated with eukaryotic hosts (Siozios et al., 2013). Different ankyrin repeat proteins have been identified that regulate host-pathogen and host-symbiont interactions (Kumagai et al., 2007; Pan et al., 2008; Nguyen et al., 2014). Aphids are well known for their interactions with associated obligate/facultative symbionts and as vectors for many plant viruses, and expression of horizontally acquired ankyrin repeat proteins was detected previously in aphid bacteriocyte and/or gut transcriptomes (Duncan et al., 2016; Feng et al., 2018). The functions of these proteins in mediating interactions between aphids and their symbionts will require further investigation. Nevertheless, our results show that these horizontally transferred ankyrin repeat proteins are important for aphid survival (Figure 4).

### HTGs in whitefly

In the whitefly genome, 64 genes were predicted to be horizontally transferred from bacteria and 78 genes were predicted to be horizontally transferred from fungi (Chen et al., 2016). Here, we have targeted five whitefly HTGs, which were chosen based on their predicted metabolic functions, using plant-mediated VIGS and demonstrated that knocking down HTGs significantly reduced whitefly survivorship (Figure 7). This suggested that HTGs could be targets for the purpose of whitefly control.

Given that horizontal transfer of functionally expressed microbial genes into insect germlines is rare on an evolutionary timescale, there is likely a selective advantage to having these genes expressed in whiteflies. This was confirmed by the observation that VIGS of both SQS and DAPDC reduced whitefly survival relative to control plants (Figure 7D). Transient expression knockdown of SQS and DAPDC, as well as other horizontally transferred genes will enable future research to study the functions of these genes in whitefly metabolism. Due to their importance for whitefly survival, as well their absence in beneficial insects such as ladybugs and lacewings, horizontally transferred genes also are attractive targets for controlling whiteflies by RNA interference.

### Avoiding negative effects in beneficial species

Despite the reduction in aphid reproduction and/or survival when HTGs were targeted by RNAi, we did not observe any negative effects on the survival of *C. septempunctata* larvae that were feeding on these aphids. This is consistent with the lack of target sequence homology in the beetles, which do not have the HTGs. However, it is also possible that there are indirect negative effects on predatory insects feeding on aphids that were targeted with RNAi constructs. For instance, silencing cathepsin L gene expression in *M. persicae* reduced their protein content and made the less suitable as prey for *C. septempunctata* (Rauf et al., 2019).

### Conclusions

The well-established *N. benthamiana* VIGS system, which allows rapid cloning of target-specific constructs (Liu et al., 2002), will allow screening of other horizontally transferred genes to identify those that are most suitable for hemipteran pest control. Such assays will be facilitated by the use of *asat2* mutant *N. benthamiana*, which improves aphid growth and allows *B. tabaci* survival (Feng et al., 2021). For the majority of tested *M. persicae* and *B. tabaci* HTGs, knocking down expression by means of plant-mediated VIGS decreased survival and/or reproduction. Importantly, no negative effects were observed on *C. septempunctata*. Together, our results indicate that HTGs have potential as efficient and biologically safe targets for aphid and whitefly population control by plant-mediated RNA interference.

## Acknowledgements

This research funded through United Stated Department of Agriculture – Agriculture and Food Research Initiative grant 2021-67013-33565 to GJ, agreement HR0011-17-2-0053 from the Defense Advanced Research Projects Agency (DARPA) Insect Allies Program with the Boyce Thompson Institute, and Binational Agricultural Research and Development Fund grant FI-471-2012 to VT.

## Author contributions

GJ, HF, and VT designed experiments. HF, VT, SS, SH, and FA: cloned genes, constructed virus vectors, measured gene expression, and conducted insect bioassays. HF, WC, and GJ analyzed data. HF and GJ wrote the manuscript. ZF, TU, and GJ obtained funding and provided other essential resources. All authors approved the final manuscript.

---

**Figure S1.**
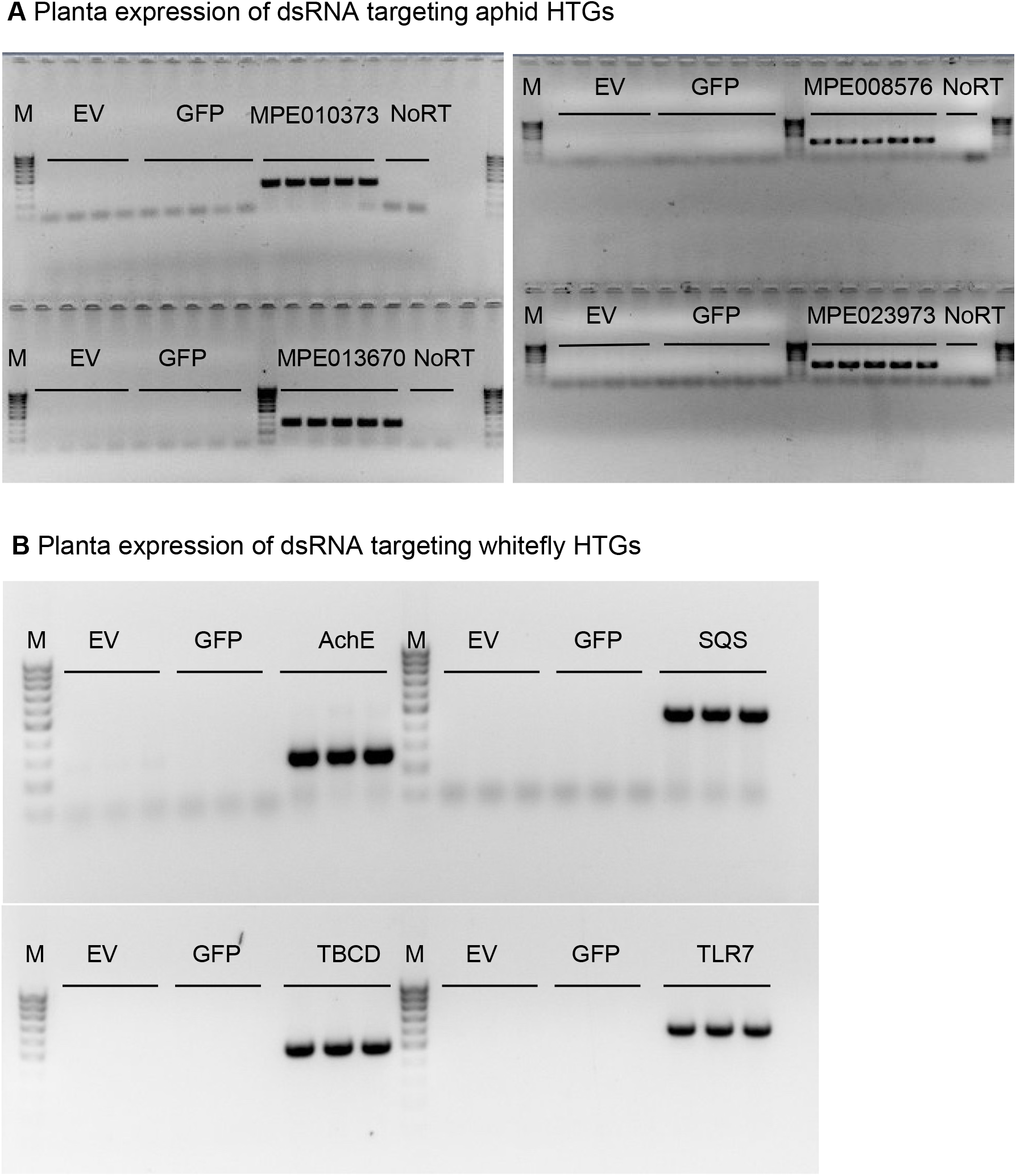
Examples of planta expression of dsRNAs that targeting horizontally transferred genes. (**A**) aphid HTG dsRNAs. (**B**) Whitefly HTG dsRNAs. EV: plants that infiltrated with TRV2 empty vector; GFP: plants that infiltrated with TRV2 that expresses a fragment targeting GFP. NoRT: no reverse transcriptase control during cDNA synthesis. M: Promega 100 bp DNA ladder.

**Figure S2.**
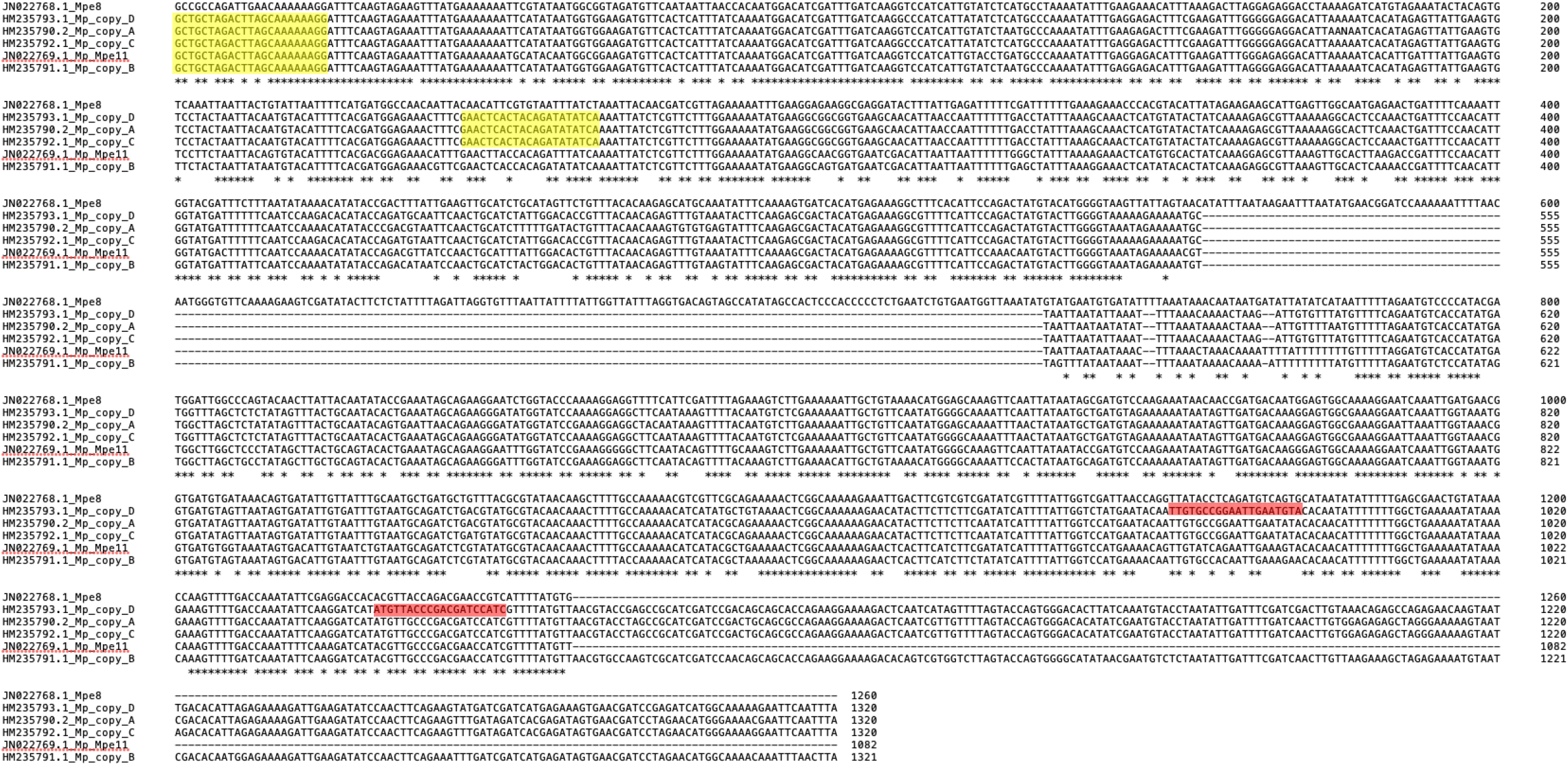
Carotenoid desaturase nucleotide alignment. Primers used for cloning TRV-*Tor* are highlighted in yellow and primers used for qPCR are highlighted in red.

**Figure S3.**
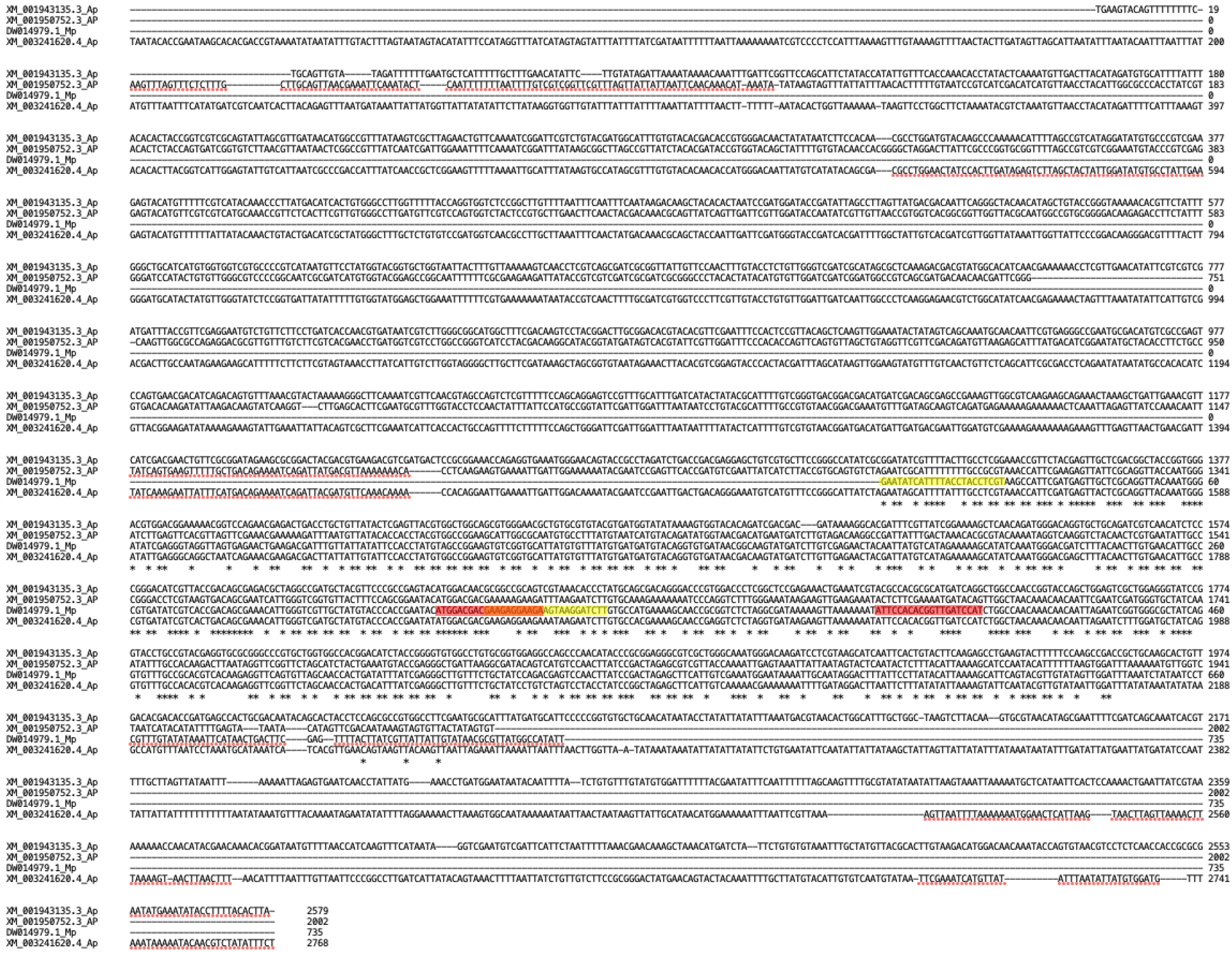
Carotenoid cyclase synthase nucleotide alignment. Primers used for cloning TRV*-CarRP* are highlighted in yellow and primers used for qPCR are highlighted in red.

**Figure S4.**
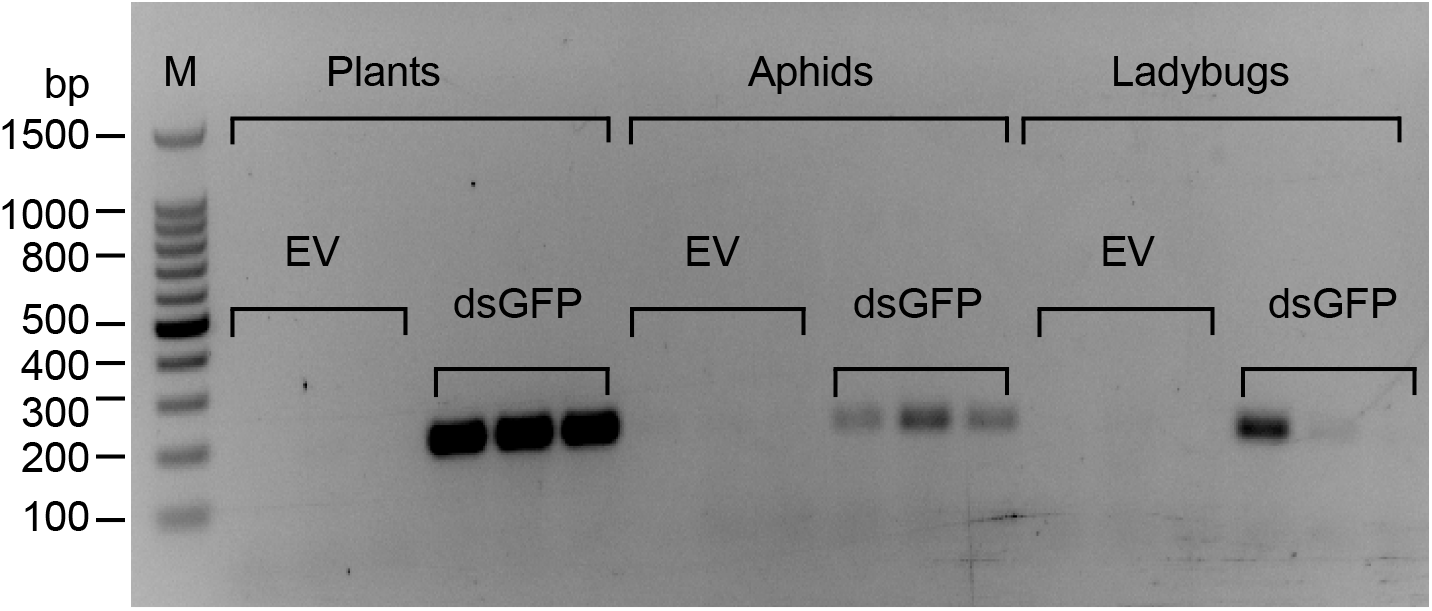
Tri-trophic persistence of dsRNA using a dsRNA targeting GFP. Agarose gel image showing the detection of dsGFP in adult *Cryptolaemus montrouzieri* ladybugs after consuming *Myzus persicae* aphids that fed on *N. benthamiana* infiltrated with TRV2-dsGFP. The expected amplicon size is about 180 bp, which is consistent to the band size shown in gel. PCR from cDNA samples of the following treatments shown in each lane: Plants: plants that infiltrated with TRV2 empty vector (EV) or TRV2-*GFP* vector (dsGFP); Aphids: aphids fed on EV and dsGFP VIGS plants; Ladybugs: ladybugs fed on aphids that were fed on EV and dsGFP VIGS plants. Each treatment was performed with three biological replicates (shown as three lanes), and this experiment was repeated twice. M: Promega 100 bp DNA ladder.

## References

Abdellatef E, Will T, Koch A, Imani J, Vilcinskas A, Kogel KH (2015) Silencing the expression of the salivary sheath protein causes transgenerational feeding suppression in the aphid Sitobion avenae. Plant Biotechnology Journal 13: 849–857

Bass C, Puinean AM, Zimmer CT, Denholm I, Field LM, Foster SP, Gutbrod O, Nauen R, Slater R, Williamson MS (2014) The evolution of insecticide resistance in the peach potato aphid, Myzus persicae. Insect Biochem Mol Biol 51: 41–51

Bhatia V, Bhattacharya R, Uniyal PL, Singh R, Niranjan RS (2012) Host generated siRNAs attenuate expression of serine protease gene in Myzus persicae. PLoS One 7: e46343

Blackman RL (1980) Chromosome-Numbers in the Aphididae and Their Taxonomic Significance. Systematic Entomology 5: 7–25

Bombarely A, Rosli HG, Vrebalov J, Moffett P, Mueller LA, Martin GB (2012) A draft genome sequence of Nicotiana benthamiana to enhance molecular plant-microbe biology research. Mol Plant Microbe Interact 25: 1523–1530

Breeden L, Nasmyth K (1987) Similarity between cell-cycle genes of budding yeast and fission yeast and the Notch gene of Drosophila. Nature 329: 651–654

Bruna T, Hoff KJ, Lomsadze A, Stanke M, Borodovsky M (2021) BRAKER2: automatic eukaryotic genome annotation with GeneMark-EP plus and AUGUSTUS supported by a protein database. Nar Genomics and Bioinformatics 3

Cantarel BL, Korf I, Robb SMC, Parra G, Ross E, Moore B, Holt C, Alvarado AS, Yandell M (2008) MAKER: An easy-to-use annotation pipeline designed for emerging model organism genomes. Genome Research 18: 188–196

Casacuberta JM, Devos Y, du Jardin P, Ramon M, Vaucheret H, Nogue F (2015) Biotechnological uses of RNAi in plants: risk assessment considerations. Trends Biotechnol 33: 145–147

Chen W, Hasegawa DK, Kaur N, Kliot A, Pinheiro PV, Luan J, Stensmyr MC, Zheng Y, Liu W, Sun H, Xu Y, Luo Y, Kruse A, Yang X, Kontsedalov S, Lebedev G, Fisher TW, Nelson DR, Hunter WB, Brown JK, Jander G, Cilia M, Douglas AE, Ghanim M, Simmons AM, Wintermantel WM, Ling KS, Fei Z (2016) The draft genome of whitefly Bemisia tabaci MEAM1, a global crop pest, provides novel insights into virus transmission, host adaptation, and insecticide resistance. BMC Biol 14: 110

Chen W, Shakir S, Bigham M, Richter A, Fei Z, Jander G (2019) Genome sequence of the corn leaf aphid (Rhopalosiphum maidis Fitch). Gigascience 8

Chin CS, Peluso P, Sedlazeck FJ, Nattestad M, Concepcion GT, Clum A, Dunn C, O’Malley R, Figueroa-Balderas R, Morales-Cruz A, Cramer GR, Delledonne M, Luo CY, Ecker JR, Cantu D, Rank DR, Schatz MC (2016) Phased diploid genome assembly with single-molecule real-time sequencing. Nature Methods 13: 1050-+

Chougule NP, Bonning BC (2012) Toxins for transgenic resistance to hemipteran pests. Toxins (Basel) 4: 405–429

Chung SH, Feng H, Jander G (2021) Engineering pest tolerance through plant-mediated RNA interference. Curr Opin Plant Biol 60: 102029

Chung SH, Jing X, Luo Y, Douglas AE (2018) Targeting symbiosis-related insect genes by RNAi in the pea aphid-Buchnera symbiosis. Insect Biochem Mol Biol 95: 55–63

Coleman AD, Wouters RH, Mugford ST, Hogenhout SA (2015) Persistence and transgenerational effect of plant-mediated RNAi in aphids. J Exp Bot 66: 541–548

Crisp A, Boschetti C, Perry M, Tunnacliffe A, Micklem G (2015) Expression of multiple horizontally acquired genes is a hallmark of both vertebrate and invertebrate genomes. Genome Biol 16: 50

Daimon T, Taguchi T, Meng Y, Katsuma S, Mita K, Shimada T (2008) Beta-fructofuranosidase genes of the silkworm, Bombyx mori: insights into enzymatic adaptation of B. mori to toxic alkaloids in mulberry latex. J Biol Chem 283: 15271–15279

Ding BY, Niu J, Shang F, Yang L, Zhang W, Smagghe G, Wang JJ (2020) Parental silencing of a horizontally transferred carotenoid desaturase gene causes a reduction of red pigment and fitness in the pea aphid. Pest Manag Sci

Dudchenko O, Batra SS, Omer AD, Nyquist SK, Hoeger M, Durand NC, Shamim MS, Machol I, Lander ES, Aiden AP, Aiden EL (2017) De novo assembly of the Aedes aegypti genome using Hi-C yields chromosome-length scaffolds. Science 356: 92–95

Duncan RP, Feng H, Nguyen DM, Wilson ACC (2016) Gene family expansions in aphids maintained by endosymbiotic and nonsymbiotic traits. Genome Biology and Evolution 8: 753–764

Durand NC, Shamim MS, Machol I, Rao SSP, Huntley MH, Lander ES, Aiden EL (2016) Juicer Provides a One-Click System for Analyzing Loop-Resolution Hi-C Experiments. Cell Systems 3: 95–98

Elzinga DA, De Vos M, Jander G (2014) Suppression of plant defenses by a Myzus persicae (green peach aphid) salivary effector protein. Mol Plant Microbe Interact 27: 747–756

Feng H, Acosta-Gamboa L, Kruse LH, Tracy JD, Chung SH, Fereira ARN, Shakir S, Xu H, Sunter G, Gore MA, Casteel CL, Moghe GD, Jander G (2021) Acylsugars protect Nicotiana benthamiana against insect herbivory and desiccation. Researchsquare

Feng H, Wang L, Wuchty S, Wilson ACC (2018) microRNA regulation in an ancient obligate endosymbiosis. Molecular Ecology 27: 1777–1793

Fernandez-Pozo N, Rosli HG, Martin GB, Mueller LA (2015) The SGN VIGS tool: user-friendly software to design virus-induced gene silencing (VIGS) constructs for functional genomics. Mol Plant 8: 486–488

Fulton M, Chunwongse J, Tanksley SD (1995) Microprep protocol for extraction of DNA from tomato and other herbaceous plants. Plant Mol Biol Rep 13: 207–209

Gotoh O (2008) Direct mapping and alignment of protein sequences onto genomic sequence. Bioinformatics 24: 2438–2444

Guindon S, Delsuc F, Dufayard JF, Gascuel O (2009) Estimating maximum likelihood phylogenies with PhyML. Methods Mol Biol 537: 113–137

Guo HY, Song XG, Wang GL, Yang K, Wang Y, Niu LB, Chen XY, Fang RX (2014) Plant-generated artificial small RNAs mediated aphid resistance. Plos One 9

Han YJ, Wessler SR (2010) MITE-Hunter: a program for discovering miniature inverted-repeat transposable elements from genomic sequences. Nucleic Acids Research 38

Hansen AK, Moran NA (2011) Aphid genome expression reveals host-symbiont cooperation in the production of amino acids. Proceedings of the National Academy of Sciences of the United States of America 108: 2849–2854

Horn T, Boutros M (2010) E-RNAi: a web application for the multi-species design of RNAi reagents--2010 update. Nucleic Acids Res 38: W332–339

Husnik F, McCutcheon JP (2018) Functional horizontal gene transfer from bacteria to eukaryotes. Nat Rev Microbiol 16: 67–79

Husnik F, Nikoh N, Koga R, Ross L, Duncan RP, Fujie M, Tanaka M, Satoh N, Bachtrog D, Wilson ACC, von Dohlen CD, Fukatsu T, McCutcheon JP (2013) Horizontal gene transfer from diverse bacteria to an insect genome enables a tripartite nested mealybug symbiosis. Cell 153: 1567–1578

Ibrahim AB, Monteiro TR, Cabral GB, Aragao FJL (2017) RNAi-mediated resistance to whitefly (Bemisia tabaci) in genetically engineered lettuce (Lactuca sativa). Transgenic Research 26: 613–624

International Aphid Genomics Consortium (2010) Genome sequence of the pea aphid Acyrthosiphon pisum. PLoS Biology 8: e1000313

Jurka J, Kapitonov VV, Pavlicek A, Klonowski P, Kohany O, Walichiewicz J (2005) Repbase update, a database of eukaryotic repetitive elements. Cytogenetic and Genome Research 110: 462–467

Kanakala S, Kontsedalov S, Lebedev G, Ghanim M (2019) Plant-Mediated Silencing of the Whitefly Bemisia tabaci Cyclophilin B and Heat Shock Protein 70 Impairs Insect Development and Virus Transmission. Front Physiol 10: 557

Kirsch R, Gramzow L, Theissen G, Siegfried BD, Ffrench-Constant RH, Heckel DG, Pauchet Y (2014) Horizontal gene transfer and functional diversification of plant cell wall degrading polygalacturonases: Key events in the evolution of herbivory in beetles. Insect Biochemistry and Molecular Biology 52: 33–50

Koch RL, Hodgson EW, Knodel JJ, Varenhorst AJ, Potter BD (2018) Management of Insecticide-Resistant Soybean Aphids in the Upper Midwest of the United States. Journal of Integrated Pest Management 9

Kumagai H, Hakoyama T, Umehara Y, Sato S, Kaneko T, Tabata S, Kouchi H (2007) A novel ankyrin-repeat membrane protein, IGN1, is required for persistence of nitrogen-fixing symbiosis in root nodules of Lotus japonicus. Plant Physiol 143: 1293–1305

Larkin MA, Blackshields G, Brown NP, Chenna R, McGettigan PA, McWilliam H, Valentin F, Wallace IM, Wilm A, Lopez R, Thompson JD, Gibson TJ, Higgins DG (2007) Clustal W and Clustal X version 2.0. Bioinformatics 23: 2947–2948

Li H, Durbin R (2009) Fast and accurate short read alignment with Burrows-Wheeler transform. Bioinformatics 25: 1754–1760

Liu YL, Schiff M, Dinesh-Kumar SP (2002) Virus-induced gene silencing in tomato. Plant Journal 31: 777–786

Livak KJ, Schmittgen TD (2001) Analysis of relative gene expression data using real-time quantitative PCR and the 2(T)(-Delta Delta C) method. Methods 25: 402–408

Lomsadze A, Burns PD, Borodovsky M (2014) Integration of mapped RNA-Seq reads into automatic training of eukaryotic gene finding algorithm. Nucleic Acids Research 42

Lu Y, Wu K, Jiang Y, Xia B, Li P, Feng H, Wyckhuys KA, Guo Y (2010) Mirid bug outbreaks in multiple crops correlated with wide-scale adoption of Bt cotton in China. Science 328: 1151–1154

Luan JB, Chen W, Hasegawa DK, Simmons AM, Wintermantel WM, Ling KS, Fei Z, Liu SS, Douglas AE (2015) Metabolic coevolution in the bacterial symbiosis of whiteflies and related plant sap-feeding insects. Genome Biology and Evolution 7: 2635–2647

Luan JB, Ghanim M, Liu SS, Czosnek H (2013) Silencing the ecdysone synthesis and signaling pathway genes disrupts nymphal development in the whitefly. Insect Biochem Mol Biol 43: 740–746

Malik HJ, Raza A, Amin I, Scheffler JA, Scheffler BE, Brown JK, Mansoor S (2016) RNAi-mediated mortality of the whitefly through transgenic expression of double-stranded RNA homologous to acetylcholinesterase and ecdysone receptor in tobacco plants. Scientific Reports 6

Mao JJ, Zeng FR (2014) Plant-mediated RNAi of a gap gene-enhanced tobacco tolerance against the Myzus persicae. Transgenic Research 23: 145–152

Mathers TC, Chen Y, Kaithakottil G, Legeai F, Mugford ST, Baa-Puyoulet P, Bretaudeau A, Clavijo B, Colella S, Collin O, Dalmay T, Derrien T, Feng H, Gabaldon T, Jordan A, Julca I, Kettles GJ, Kowitwanich K, Lavenier D, Lenzi P, Lopez-Gomollon S, Loska D, Mapleson D, Maumus F, Moxon S, Price DR, Sugio A, van Munster M, Uzest M, Waite D, Jander G, Tagu D, Wilson ACC, van Oosterhout C, Swarbreck D, Hogenhout SA (2017) Rapid transcriptional plasticity of duplicated gene clusters enables a clonally reproducing aphid to colonise diverse plant species. Genome Biology 18: 27

Mendelsohn M, Kough J, Vaituzis Z, Matthews K (2003) Are Bt crops safe? Nat Biotechnol 21: 1003–1009

Moran NA, Jarvik T (2010) Lateral transfer of genes from fungi underlies carotenoid production in aphids. Science 328: 624–627

Mount DW (2007) Using the Basic Local Alignment Search Tool (BLAST). CSH Protoc 2007: pdb top17

Mutti NS, Park Y, Reese JC, Reeck GR (2006) RNAi knockdown of a salivary transcript leading to lethality in the pea aphid, Acyrthosiphon pisum. J Insect Sci 6: 1–7

Nguyen MTHD, Liu M, Thomas T (2014) Ankyrin-repeat proteins from sponge symbionts modulate amoebal phagocytosis. Molecular Ecology 23: 1635–1645

Nikoh N, McCutcheon JP, Kudo T, Miyagishima SY, Moran NA, Nakabachi A (2010) Bacterial genes in the aphid genome: absence of functional gene transfer from Buchnera to its host. PLoS Genet 6: e1000827

Nikoh N, Nakabachi A (2009) Aphids acquired symbiotic genes via lateral gene transfer. BMC Biol 7: 12

Novakova E, Moran NA (2012) Diversification of genes for carotenoid biosynthesis in aphids following an ancient transfer from a fungus. Mol Biol Evol 29: 313–323

Pan X, Luhrmann A, Satoh A, Laskowski-Arce MA, Roy CR (2008) Ankyrin repeat proteins comprise a diverse family of bacterial type IV effectors. Science 320: 1651–1654

Parker BJ, Brisson JA (2019) A laterally transferred viral gene modifies aphid wing plasticity. Curr Biol 29: 2098–2103 e2095

Pauchet Y, Heckel DG (2013) The genome of the mustard leaf beetle encodes two active xylanases originally acquired from bacteria through horizontal gene transfer. Proc. Biol. Sci. 280: 20131021

Pauchet Y, Kirsch R, Giraud S, Vogel H, Heckel DG (2014) Identification and characterization of plant cell wall degrading enzymes from three glycoside hydrolase families in the cerambycid beetle Apriona japonica. Insect Biochemistry and Molecular Biology 49: 1–13

Pessoa R, Rossi GD, Busoli AC (2016) Transgenic Cotton-Fed Bemisia tabaci (Gennadius) (Hemiptera: Aleyrodidae) Affects the Parasitoid Encarsia desantisi Viggiani (Hymenoptera: Aphelinidae) Development. Neotrop Entomol 45: 102–106

Pitino M, Coleman AD, Maffei ME, Ridout CJ, Hogenhout SA (2011) Silencing of aphid genes by dsRNA feeding from plants. Plos One 6

Ramsey JS, Elzinga DA, Sarkar P, Xin YR, Ghanim M, Jander G (2014) Adaptation to nicotine feeding in Myzus persicae. J Chem Ecol 40: 869–877

Ramsey JS, Wilson AC, De Vos M, Sun Q, Tamborindeguy C, Winfield A, Malloch G, Smith DM, Fenton B, Gray SM, Jander G (2007) Genomic resources for Myzus persicae: EST sequencing, SNP identification, and microarray design. BMC Genomics 8: 423

Rauf I, Asif M, Amin I, Naqvi RZ, Umer N, Mansoor S, Jander G (2019) Silencing cathepsin L expression reduces Myzus persicae protein content and the nutritional value as prey for Coccinella septempunctata. Insect Mol Biol 28: 785–797

Raza A, Malik HJ, Shafiq M, Amin I, Scheffler JA, Scheffler BE, Mansoor S (2016) RNA Interference based Approach to Down Regulate Osmoregulators of Whitefly (Bemisia tabaci): Potential Technology for the Control of Whitefly. PLoS One 11: e0153883

Rodrigues TB, Mishra SK, Sridharan K, Barnes ER, Alyokhin A, Tuttle R, Kokulapalan W, Garby D, Skizim NJ, Tang YW, Manley B, Aulisa L, Flannagan RD, Cobb C, Narva KE (2021) First Sprayable Double-Stranded RNA-Based Biopesticide Product Targets Proteasome Subunit Beta Type-5 in Colorado Potato Beetle (Leptinotarsa decemlineata). Front Plant Sci 12: 728652

RStudio Team (2020) RStudio: Integrated Development for R. In http://www.rstudio.com/. RStudio, Boston, MA

Sears MK, Hellmich RL, Stanley-Horn DE, Oberhauser KS, Pleasants JM, Mattila HR, Siegfried BD, Dively GP (2001) Impact of Bt corn pollen on monarch butterfly populations: a risk assessment. Proc Natl Acad Sci U S A 98: 11937–11942

Sedgwick SG, Smerdon SJ (1999) The ankyrin repeat: a diversity of interactions on a common structural framework. Trends Biochem Sci 24: 311–316

Senthil-Kumar M, Mysore KS (2014) Tobacco rattle virus-based virus-induced gene silencing in Nicotiana benthamiana. Nat Protoc 9: 1549–1562

Simao FA, Waterhouse RM, Ioannidis P, Kriventseva EV, Zdobnov EM (2015) BUSCO: Assessing genome assembly and annotation completeness with single-copy orthologs. Bioinformatics 31: 3210–3212

Siozios S, Ioannidis P, Klasson L, Andersson SG, Braig HR, Bourtzis K (2013) The diversity and evolution of Wolbachia ankyrin repeat domain genes. PLoS One 8: e55390

Sloan DB, Nakabachi A, Richards S, Qu J, Murali SC, Gibbs RA, Moran NA (2014) Parallel histories of horizontal gene transfer facilitated extreme reduction of endosymbiont genomes in sap-feeding insects. Molecular Biology and Evolution 31: 857–871

Smith TE, Li Y, Perreau J, Moran NA (2022) Elucidation of host and symbiont contributions to peptidoglycan metabolism based on comparative genomics of eight aphid subfamilies and their Buchnera. PLoS Genet 18: e1010195

Soucy SM, Huang J, Gogarten JP (2015) Horizontal gene transfer: building the web of life. Nat Rev Genet 16: 472–482

Stanke M, Waack S (2003) Gene prediction with a hidden Markov model and a new intron submodel. Bioinformatics 19: Ii215–Ii225

Sujii ER, Togni PH, de ARP, de ABT, Milane PV, Paula DP, Pires CS, Fontes EM (2013) Field evaluation of Bt cotton crop impact on nontarget pests: cotton aphid and boll weevil. Neotrop Entomol 42: 102–111

Thakur N, Upadhyay SK, Verma PC, Chandrashekar K, Tuli R, Singh PK (2014) Enhanced Whitefly Resistance in Transgenic Tobacco Plants Expressing Double Stranded RNA of v-ATPase A Gene. Plos One 9

Therneau TM, Grambsch PM (2000) Modeling surivival data: extending the cox model. Springer-Verlag

Tian JC, Yao J, Long LP, Romeis J, Shelton AM (2015) Bt crops benefit natural enemies to control non-target pests. Sci Rep 5: 16636

Tzin V, Yang X, Jing X, Zhang K, Jander G, Douglas AE (2015) RNA interference against gut osmoregulatory genes in phloem-feeding insects. J Insect Physiol 79: 105–112

Vaghchhipawala Z, Rojas CM, Senthil-Kumar M, Mysore KS (2011) Agroinoculation and agroinfiltration: simple tools for complex gene function analyses. Methods Mol Biol 678: 65–76

Velasquez AC, Chakravarthy S, Martin GB (2009) Virus-induced gene silencing (VIGS) in Nicotiana benthamiana and tomato. J Vis Exp

Verster KI, Wisecaver JH, Karageorgi M, Duncan RP, Gloss AD, Armstrong EE, Price DK, Menon AR, Ali ZM, Whiteman NK (2019) Horizontal Transfer of Bacterial Cytolethal Distending Toxin B Genes to Insects. Mol Biol Evol 36: 2105–2110

Walker BJ, Abeel T, Shea T, Priest M, Abouelliel A, Sakthikumar S, Cuomo CA, Zeng QD, Wortman J, Young SK, Earl AM (2014) Pilon: An Integrated Tool for Comprehensive Microbial Variant Detection and Genome Assembly Improvement. Plos One 9

Wybouw N, Pauchet Y, Heckel DG, Van Leeuwen T (2016) Horizontal gene transfer contributes to the evolution of arthropod herbivory. Genome Biol. Evol. 8: 1785–1801

Xia J, Guo Z, Yang Z, Han H, Wang S, Xu H, Yang X, Yang F, Wu Q, Xie W, Zhou X, Dermauw W, Turlings TCJ, Zhang Y (2021) Whitefly hijacks a plant detoxification gene that neutralizes plant toxins. Cell 184: 1693–1705 e1617

Xie X, Shang F, Ding BY, Yang L, Wang JJ (2022) Assessment of a zinc finger protein gene (MPZC3H10) as potential RNAi target for green peach aphid Myzus persicae control. Pest Manag Sci 78: 4956–4962

Xu L, Duan X, Lv Y, Zhang X, Nie Z, Xie C, Ni Z, Liang R (2014) Silencing of an aphid carboxylesterase gene by use of plant-mediated RNAi impairs Sitobion avenae tolerance of Phoxim insecticides. Transgenic Res 23: 389–396

Zhao C, Doucet D, Mittapalli O (2014) Characterization of horizontally transferred beta-fructofuranosidase (ScrB) genes in Agrilus planipennis. Insect Mol Biol 23: 821–832

Zhong S, Joung JG, Zheng Y, Chen YR, Liu B, Shao Y, Xiang JZ, Fei Z, Giovannoni JJ (2011) High-throughput Illumina strand-specific RNA sequencing library preparation. Cold Spring Harb Protoc 2011: 940–949

